# An ultra-low background far-red light-responsive optogenetic tool based on an engineered biliverdin-binding domain

**DOI:** 10.1101/2025.11.29.691301

**Authors:** Giang N. T. Le, Lam M. T. Pham, Boyan Xue, Maruti Uppalapati, G. Andrew Woolley

## Abstract

The robustness and broad applicability of an optogenetic tool depends heavily on the properties of the underlying photoreceptor protein and its cognate binding partner - the light responsive ‘core’. Current red light optogenetic systems for use in mammalian cells all rely on phytochrome based photoreceptors. These are large (70 kDa) proteins that act as dimers, thereby enforce dimerization on attached proteins. Naturally occurring or engineered binding partners can function effectively in certain cases, but large size, complex mode of interaction, background binding, relatively weak affinity and/or low fold changes between on and off states are significant limitations. Using structure-based design and directed evolution we developed a small (17 kDa) monomeric bilverdin binding photoreceptor FenixS, and a highly selective, high-affinity binder, Ash1 (6 kDa). Negligible off-state binding and a >1200-fold increase in binding affinity upon 700 nm illumination result in a high performance, ultra-low background, light responsive core for a diverse range of applications. An optogenetic tool for red light activation of transcription in mammalian cells based on the FenixS-Ash1 core exhibits robust performance without the need for biliverdin supplementation.

## Introduction

Optogenetic tools offer researchers a powerful technology for controlling protein-protein interactions (PPIs), enabling manipulation of cellular processes with high spatiotemporal resolution. The development of these tools has enabled groundbreaking research on the roles of dynamic gene expression^1,2^, the functions of biomolecular condensates^3^, and the spatiotemporal coordination of morphogens during embryogenesis^4^. While blue light-responsive systems such as CRY2-CIB1^5^, LOVTRAP^6^, iLID^7^, and Magnet^8^, have been widely adopted over the past decade, red and near-infrared (NIR) light-responsive tools have gained increasing attention and have become a focal point for development^9,10^ due to the superior tissue penetration and reduced phototoxicity of red and NIR light^11^. At the core of all these heterodimerizing optogenetic systems is a photoswitchable protein that undergoes a conformational change in response to a specific wavelength of light and a binder that selectively interacts with one conformational state of the photoswitchable protein.

The core photoswitchable components of current red-light responsive tools have significant limitations. First, except for the BICYCL system which uses the non-native phycocyanobilin chromophore^12^, the photoreceptors used in all current red-light systems are either plant or bacterial phytochromes. These are large (∼70 kDa) proteins which limit their use in viral (AAV) delivery systems, and dimeric^13,14^ which leads to potentially unwanted dimerization of their fusion partners. Second, while bacterial phytochromes use biliverdin as a chromophore, which is present in mammalian cells, their binding partners are either large (PpsR2), lead to background activity due to apo-protein binding (Q-PAS1), have relatively weak affinity (*K*_d_ = 470 nM; nanoRED^15^), or limited (50-fold) state selectivity (MagRed^16^). While non-linear responses in biological systems can mean that relatively small fold-changes in protein-protein interaction affinity nevertheless have functional consequences, a large intrinsic fold change and low basal activity in the core photoswitchable components are essential for a robust optogenetic tools suitable for diverse applications^17,18^.

Here, we report the development of a small, monomeric, red light-activable optogenetic system based on a BV-binding photoswitchable protein, FenixS (17 kDa), and a highly selective, high-affinity binder, Ash1 (6 kDa). FenixS was generated through structure-guided mutagenesis and extensive directed evolution of miRFP670nano, resulting in a photoswitchable protein that can function in mammalian cells without the need for exogenous BV supplementation. Ash1 is a *de novo* synthetic binder derived via phage display from the albumin-binding domain of protein G (GA domain). In addition to being small and monomeric, FenixS binds Ash1 with high affinity (*K*_d_ = 42 nM) and exhibits >1,200-fold difference in binding affinity between light and dark states. The FenixS-Ash1 pair thus provides an ultra-low background activity red light-responsive core for optogenetic tool development. We demonstrate the performance and versatility of the FenixS-Ash1 pair in regulating subcellular protein localization and transcriptional activation. Furthermore, by replacing the iLID-SspB core in Corelets^19^ with FenixS-Ash1, we established a red light-inducible biomimetic system for condensate formation, enabling precise spatiotemporal control using red light. Across experimental setups and cell lines, the FenixS-Ash1 core consistently exhibited minimal background activity, with or without exogenous BV supplementation.

## Results

### Development and characterization of a BV-binding photoswitchable protein: FenixS

Naturally occurring cyanobacterial GAF domains are attractive candidates for optogenetic applications due to their compact size and spectral diversity^20^. However, one key limitation is their suboptimal BV-binding ability^12^. While some CBCR GAF domains can bind BV due to its structural similarity to PCB, it appears there has been no evolutionary pressure to optimize this interaction. Attempts to develop BV-binding CBCR-based optogenetic tools, such as BICYCL-biliverdin^12^, have not yielded a high-performance tool for use in mammalian cells. Efforts to enhance BV binding through site-directed mutagenesis (e.g., AnPixJg2_BV4^21^ and AM1_1186g2 KCAP_QV^22^) have been limited to a small set of mutations. Even AM1_C0023g2 S334G^23^, a benchmark CBCR in those studies, exhibits relatively weak BV-binding efficiency (*vide infra*)^12^. Verkhusha and colleagues used extensive directed evolution to convert the CBCR GAF domain NpR3784 into the miRFP670nano series^24–27^, proteins that bound BV well enough to allow robust detection of fluorescence in mammalian cells. However, over the course of this directed evolution campaign, photoswitchability was lost.

To develop a BV-binding photoswitchable protein based on a cyanobacteriochrome GAF protein, we selected miRFP670nano^24^ as a starting template. Our goal was to restore photoswitchability while preserving BV-binding capability. In this protein family, photoisomerization occurs via cis-trans isomerization in C15=C16 double bond, which rotates the D-ring out of the conjugated π electron system of the chromophore^28^ (Supplementary Fig. 1). We hypothesized that mutations in the proximity of the D-ring introduced during the evolution of miRFP670nano enhanced fluorescence by restricting D-ring mobility (Supplementary Fig. 2). To test this, we reverted five residues (C66F, V67M, F68Y, Y128H, and H158N; NpR3784 numbering) to their original identities in NpR3784. The resulting variant, designated Fenix R0, exhibited fluorescence that was two orders of magnitude dimmer than miRFP670nano but showed a substantial reversible change in fluorescence upon alternating irradiation with 660 nm and 530 nm light (F_ON_/F_OFF_ = 3.6; Extended Data Fig. 1). The name Fenix – a variant of “phoenix”, often associated with red colors^29^ – was chosen to reflect both the near-infrared fluorescence of the protein and its photoswitchable nature, reminiscent of the phoenix life cycle: cycling between a bright, fluorescent dark state and a non-fluorescent photo-state. Fenix R0 served as the starting point for further engineering via directed evolution.

During directed evolution, we used whole-cell fluorescence (F_ON_) and change in fluorescence intensity upon irradiation (F_ON_/F_OFF_) as proxies for BV-binding efficiency and photoswitchability, respectively. These metrics enabled screening of tens of thousands of bacterial colonies per round. The selection strategy was designed to find variants with high ON-state fluorescence, low OFF-state fluorescence, fast switching kinetics, strong fatigue resistance, and a rapid BV-binding (Fig. 1a). After 18 rounds of random and site-saturation mutagenesis (Fig. 1b,c and Extended Data Fig. 2), we obtained variant Fenix R18, which exhibited a fully reversible F_ON_/F_OFF_ ratio of 12, over at least two switching cycles under the screening conditions (i.e., crude protein extract in B-PER solution) (Supplementary Fig. 3). Compared to miRFP670nano, Fenix R18 contains 25 point-mutations (NpR3784 numbering): N44Y, L45Q, T51D, T54A, E55K, F59I, V62A, C66V, V67M, V77E, D85G, I88S, K92G, V95C, R96S, T103S, S109A, H110R, T125S, Y128H, T133A, I140V, T168S, V180A, S188I (Fig. 1c).

**Fig. 1:**
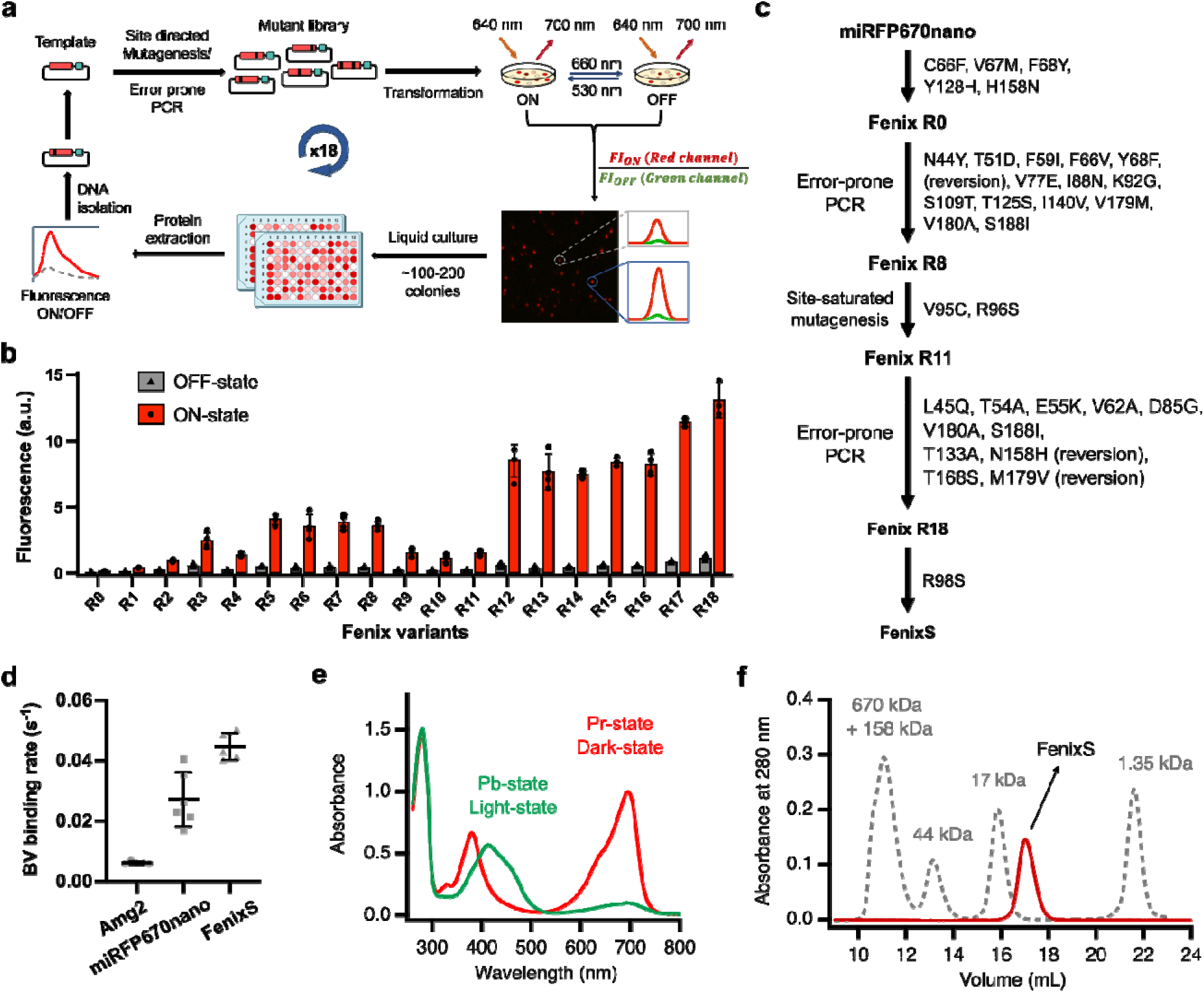
Development and characterization of FenixS. **a,** The directed evolution workflow for FenixS. Starting from the template Fenix R0, the full-length gene (red) was randomly mutated by error-prone PCR or site-saturation mutagenesis. The resulting library was ligated into a plasmid containing heme-oxygenase (green) and the product was used to transform *E. coli*. Fluorescence images of the plates after red-light (660 nm) illumination (OFF-state, red channel) and green-light (530 nm) illumination (ON-state, green channel) were overlayed. Bright (high F_ON_) and red (large F_ON_/F_OFF_) colonies were picked and cultured. Winners of each round were selected based on their brightness (F_ON_), fold-change (F_ON_/F_OFF_), and reversibility after 2 cycles (measured using crude protein extracts). Winners were used as the templates for the next round of evolution. **b,** Fluorescence of the ON and OFF states of selected variants after each round of directed evolution. The OFF-state was produced with 660 nm illumination, The ON-state was produced with 530 nm illumination. Fluorescence of both states was detected at 670 nm with excitation at 647 nm. Measurements were performed using crude protein extracts from *E. coli*. n = 3 biological replicates, mean ± sd. **c,** Lineage of FenixS variants starting from miRFP670nano. **d,** Assembly rates of apoproteins Amg2, miRFP670nano, and FenixS with BV (ratio 1.5 to 1) *in vitro*. **e,** UV-Vis absorbance spectra of the FenixS protein in its Pr (dark)-state and Pb (photo)-state. The Pb-state was obtained by irradiation for 4 minutes with 700 nm light (14 mW cm^-2^). **f,** Size-exclusion chromatography of FenixS at a concentration of 50 µM. Molecular weight standards are indicated.

Although fluorescence was used as a screening metric, it is not essential for photoswitchable proteins; indeed, fluorescence and photoswitching are two competing pathways for de-excitation of the excited state. During the directed evolution campaign, we identified a single colony with markedly darker green color suggesting efficient chromophore incorporation; this colony also had significantly reduced fluorescence (Supplementary Fig. 4). Sequencing revealed a single point mutation, R98S, located near the canonical Cys that covalently attaches to the chromophore (Supplementary Fig. 5). Interestingly, the reverse mutation (S98R) was one of the key changes introduced during the evolution of miRFP670nano from NpR3784. We introduced the R98S mutation to Fenix R18 to generate FenixS (Fig. 1c, Supplementary Fig. 6), hypothesizing that it would enhance chromophore incorporation efficiency. Indeed, FenixS exhibited a higher BV-binding rate than miRFP670nano (Fig. 1d). BV-binding rate has been shown to correlate with the holo/apo-protein ratio in mammalian expression systems^30,31^, indicating that FenixS surpasses the BV-binding efficiency threshold originally set by miRFP670nano at the outset of our directed evolution campaign. Furthermore, FenixS exhibited a sevenfold faster BV-binding rate compared to AM1_C0023g2 S334G, the benchmark natural CBCR-based photoswitchable protein noted for its efficient BV incorporation^23^.

In the dark-state, FenixS exhibits absorbance peaks at 696 nm and 380 nm which correspond to Q- and Soret-bands^32^, respectively. Upon photoconversion, there is a suppression of the Q-band and an apparent red-shift of the Soret-band to produce a blue-absorbing state (Fig. 1e). Similar spectral behavior was observed with the NpF2164g3 CBCR-GAF domain and was attributed to the addition of second Cys to C10 of chromophore during photoconversion^33^. This addition breaks π-conjugation of the chromophore leading to absorbance in the blue region of the spectrum. In the case of FenixS, the newly introduced Cys95 to may also add to C10 of the chromophore (Supplementary Fig. 1 and 5) upon photoconversion. FenixS fully recovers at 37°C in the dark and exhibits bi-exponential dark reversion with half-lives of 12 and 109 minutes for the fast and slow processes, respectively (Extended Data Fig. 3). Note that high intensity blue light can induce photobleaching due to overlap with the Soret band of the chromophore (Extended Data Fig. 4a). Photobleaching effect can be minimized by using 505-530 nm light for reverse switching (Extended Data Fig. 4b,c), as employed during the directed evolution of FenixS. Size-exclusion chromatography confirmed that FenixS is a monomer at concentrations up to 50 µM (Fig. 1f). Interestingly, FenixS elutes with a longer retention time than expected for an 18.6 kDa monomeric protein, suggesting a compact protein conformation.

### FenixS-Ash development and characterization

To develop a red light-activable optogenetic tool based on FenixS, we employed an optimized phage display-based selection strategy to generate a binder specific to the photo-state (Fig. 2a)^34,35^. In brief, twelve residues within the second and third helices of the GA scaffold were randomized to construct a naïve library, which was displayed on the pVIII coat protein of M13 phage (Fig. 2b) ^36^. After three rounds of selection and single clone ELISA screening, we identified a photo-state-specific binder, Ash0.1, which exhibits ∼30-fold selectivity for the photo-state of FenixS over the dark-state and the apo-state in phage ELISA assays (Supplementary Fig. 7a). ELISA using purified proteins confirmed the selectivity of Ash0.1 for the ^15E^Pb photo-state and gave a dissociation constant (*K*_d_) in the micromolar range (Supplementary Fig. 7b). To improve affinity, we constructed an affinity maturation library based on Ash0.1 and displayed it on the pIII coat protein (Fig. 2b). After three rounds of selection, single clone phage ELISA screening revealed enrichment of photo-state-specific binders (Supplementary Fig. 8). The six variants with the highest relative phage binding signals (Fig. 2c, Supplementary Fig. 9) were expressed, purified, and their binding affinities toward *^15Z^*Pr-, *^15E^*Pb- and apo-FenixS were determined (Fig. 2d, Extended Data Fig. 5). While all variants demonstrated selectivity for *^15E^*Pb photo-state, their binding affinities did not directly correlate with phage binding signals (Fig. 2c). For instance, Ash0.5 exhibited the highest phage binding signal but had a *K*_d_ of 1.25 µM, whereas Ash1, with a *K*_d_ of 41 nM, ranked third in phage ELISA (Fig. 2c,d and Extended Data Fig. 5). This discrepancy may be attributed to differences in display levels or phage titers (not normalized during the ELISA protocol) among variants. The three top variants – Ash1, Ash0.9, and Ash0.8 – exhibited negligible binding to *^15Z^*Pr- and apo-FenixS at the highest concentration tested (Fig. 2d, Extended Data Fig. 5). Among them, Ash1 exhibited the highest affinity (*K*_d_ = 41 nM), followed by Ash0.9 and Ash0.8 with *K*_d_ values of 310 nM and 331 nM, respectively (Fig. 2d). Size exclusion chromatography (SEC) confirmed that all three proteins were monomeric (Supplementary Fig. 10), and circular dichroism spectra indicated that they adopted a three-helix structure like the wild-type GA domain^37^ (Supplementary Fig. 11).

**Fig. 2:**
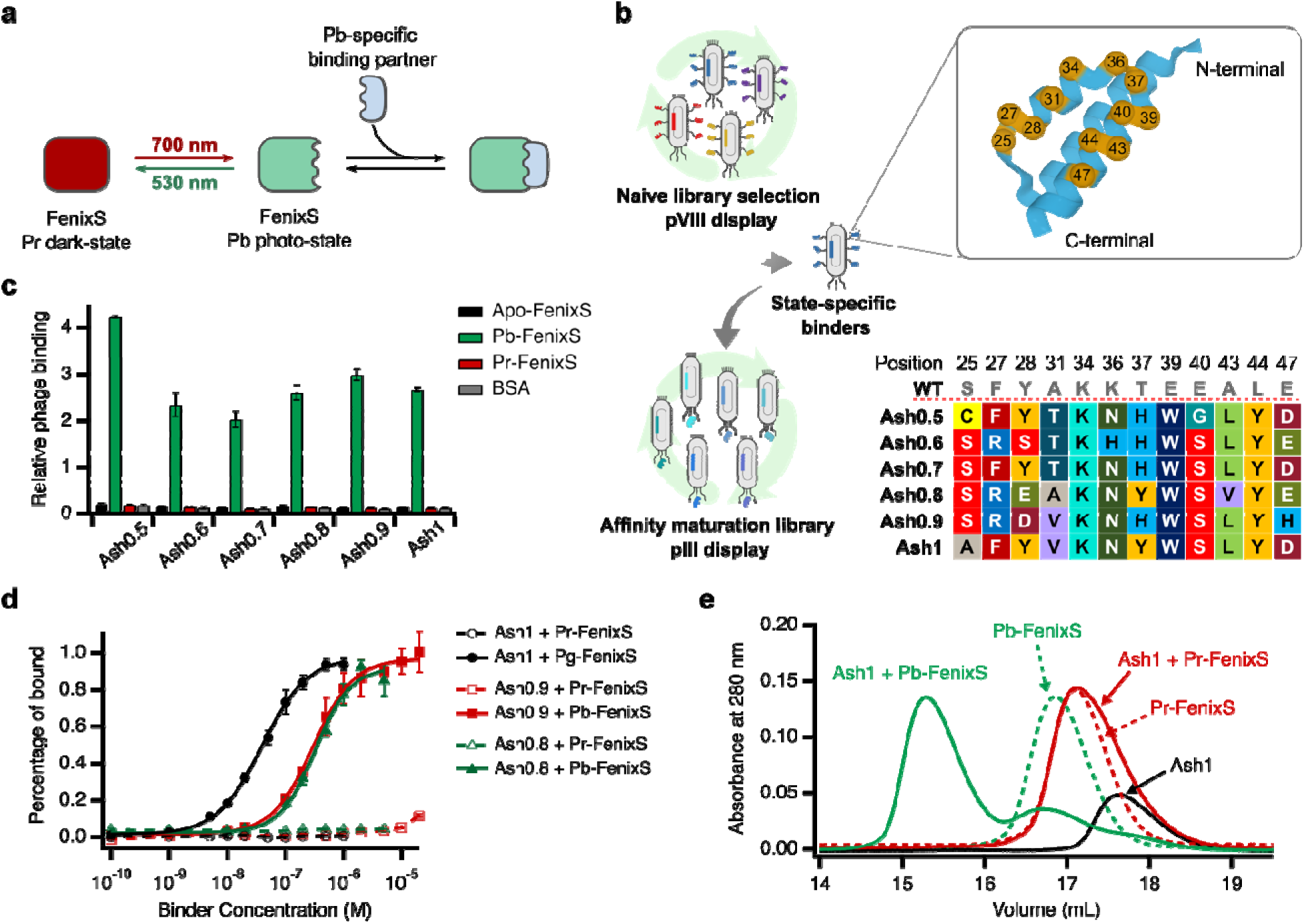
Development and characterization of the FenixS-Ashes optogenetic system. **a,** Schematic representation of FenixS and its photo-state-specific binding partner. **b,** Overview of the phage display-based selection strategy for Ashes (left). The positions of the 12 randomized residues within the second and third helices of the GA domain are indicated, along with the sequences of the photo-state-specific binders (right). Amino acid residues are highlighted with background color corresponding to their amino acid identities. Full sequences of selected binders are provided in Supplementary Fig. 9. **c,** Phage ELISA results for the six variants with the highest relative binding signals. n = 4 technical replicates, mean ± sd. Data represents results from two independent experiments. **d,** ELISAs using purified, biotinylated FenixS with purified binders Ash1, Ash0.9, and Ash0.8. Apparent dissociation constants (*K*_d_) were estimated by fitting the data to a Hill equation with a Hill slope of 1.0. n = 4 technical replicates, mean ± sd. **e,** Size exclusion chromatography (SEC) analysis of light-dependent complex formation. Free Ash1, FenixS and FenixS/Ash1 mixtures were analyzed under dark or red-light conditions. Light-adapted samples were pre-irradiated with 700 nm red LED for 5 minutes prior to loading into the column. During SEC, the column was either kept in the dark or irradiated with a 730 nm red LED array. Protein elution was monitored by absorbance at 280 nm. Shorter retention times indicate larger apparent molecular size. Note that a small fraction of Pb-FenixS remains unbound to Ash1; see Supplementary Fig. 12 for further discussion.

To assess the interaction between FenixS and Ash1 at higher concentrations, mixtures of FenixS (50 µM) and Ash1 (50 µM) were analyzed by size exclusion chromatography (SEC) under red light and in the dark (Fig. 2e, Supplementary Fig. 12). FenixS and Ash1 alone were chromatographed under identical conditions for comparison (Fig. 2e, Supplementary Fig. 12). A shift in retention time under red light (Fig. 2e) indicated formation of a complex between *^15E^*Pb-FenixS and Ash1. In contrast, no shift was observed in the dark, implying a dissociation constant (*K*_d_) > 50 µM for *^1ZE^*Pr-FenixS and Ash1. Together, ELISA and SEC data show that Ash1 exhibits over 1,200-fold selectivity for the Pb-state over Pr-state.

### Photoswitching of FenixS-Ash on agarose beads

Many biotechnological applications of optogenetic protein-protein tools occur at surfaces, such as in light-control affinity chromatography and cell surface receptor binding^38,39^. To investigate whether FenixS-Ashes interactions could be regulated on the surface of agarose beads we immobilized FenixS (or IgG as a negative control) and placed the beads in a solution containing 500 nM of mCherry-Ash1 (Fig. 3a). FenixS-coated beads were distinguished from IgG-coated beads by their far-red fluorescence (Fig. 3b). We observed red light-dependent surface localization of Ash1 on FenixS-coated beads, as indicated by mCherry fluorescence (Fig. 3c, Movie S1). Blue light, in turn, caused dissociation of the mCherry signal.

**Fig. 3:**
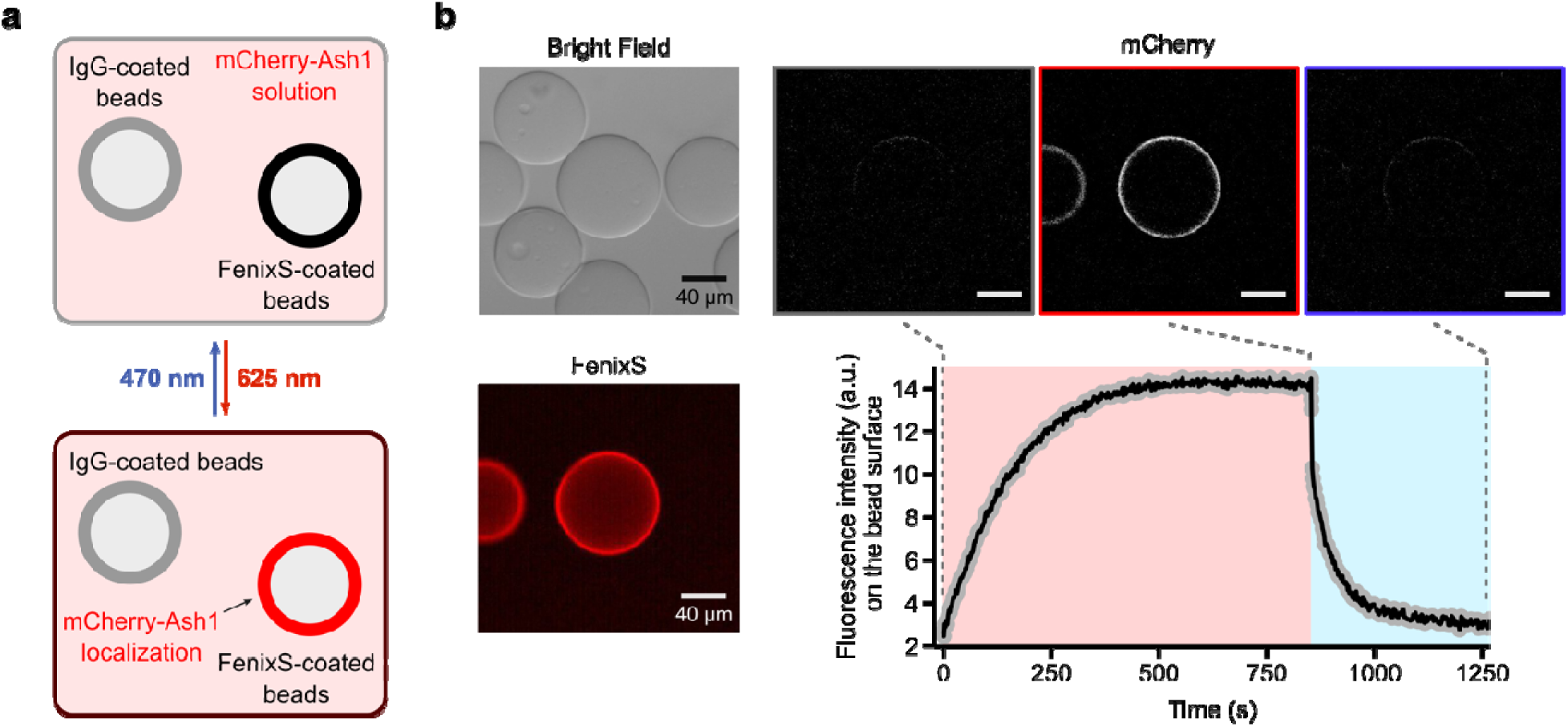
FenixS/Ash-mediated optogenetic control of protein recruitment to bead surfaces. **a,** Schematic illustration of the light-controlled recruitment of mCherry-Ash1 to FenixS-coated beads. Streptavidin-coated agarose beads were separately functionalized with biotinylated FenixS or IgG (negative control). The bead mixture was incubated in an mCherry-Ash1 solution. Upon red-light illumination, FenixS undergoes a conformational change to its photo-state, enabling the recruitment of mCherry-Ash1 to the surface of FenixS-coated beads. Blue-light illumination reverses this process by converting FenixS to its dark-state, allowing mCherry-Ash1 to diffuse away. **b,** Left panel: Brightfield and FenixS fluorescence images showing the position of IgG- and FenixS-coated beads. Beads were incubated in 500 nM mCherry-Ash1. Top-right panel: mCherry fluorescence images at indicated time points. Bottom-right panel: Quantification of mCherry fluorescence intensity on bead surfaces during alternating red (625 nm and blue (470 nm) light illumination (indicated by red and blue shading, respectively). Images are representative of four beads from two independent experiments.

### Transcriptional activation optogenetic system

Optogenetic control of gene expression represents a major application of light-responsive systems^40,41^. To assess the suitability of the FenixS-Ash1 core for light-inducible transcriptional activation, we designed constructs based on our previous work with BICYCL^12,42^ (Fig. 4a). Two constructs were generated: a FenixS-VP16 fusion and a FenixS-VP16 fusion that was also linked to the intrinsically disordered region of the human oncogene FUS_N_, previously reported to enhance transcriptional activity^43^. The performance of the system was evaluated via transient transfection into Chinese hamster ovary (CHO-K1) cell lines using a secreted alkaline phosphatase (SEAP) reporter. For initial tests we supplemented the cells with 40 uM BV to ensure FenixS was maximally chromophorylated. Under these conditions, both constructs exhibited minimal basal activity and showed >200-fold induction of SEAP expression upon red light illumination (Fig. 4b). Notably, the second design (FenixS-FUS_N_-VP16-IRES-E-Ash1) achieved 50% of the SEAP expression level of the positive control construct which signifies maximal transcriptional activation (E-VP16), with signal continuing to increase over 48 hours (Fig. 4b and Extended Data Fig. 6). We then tested if exogenous BV was required for red light activation of SEAP expression. Chromophore titration experiments (Fig. 4c) identified 20 µM biliverdin (BV) as the optimal concentration, however, 47% of maximal FenixS transcriptional activation was achieved *without* BV supplementation, indicating that endogenous BV levels are sufficient to enable holo-protein to form and function effectively in this system. To directly examine the holo/apo ratio of FenixS in mammalian cells, we fused FenixS to mCherry and expressed this construct in Hela cells. By comparing the intrinsic far-red fluorescence of FenixS (holo-form) to mCherry fluorescence (total protein), we found that the fraction of holo FenixS could be increased by adding 5 µM or 40 µM BV by ∼10-fold and 27-fold, respectively (Supplementary Fig. 13). Thus, signal amplitudes can be increased by increasing holo-protein abundance – a property shared by all phytochrome-based fluorescence proteins^44^. Importantly, although apo-FenixS remains present without BV supplementation, it acts as a bystander only, since Ash1 does not interact with the apo-form.

**Fig. 4.**
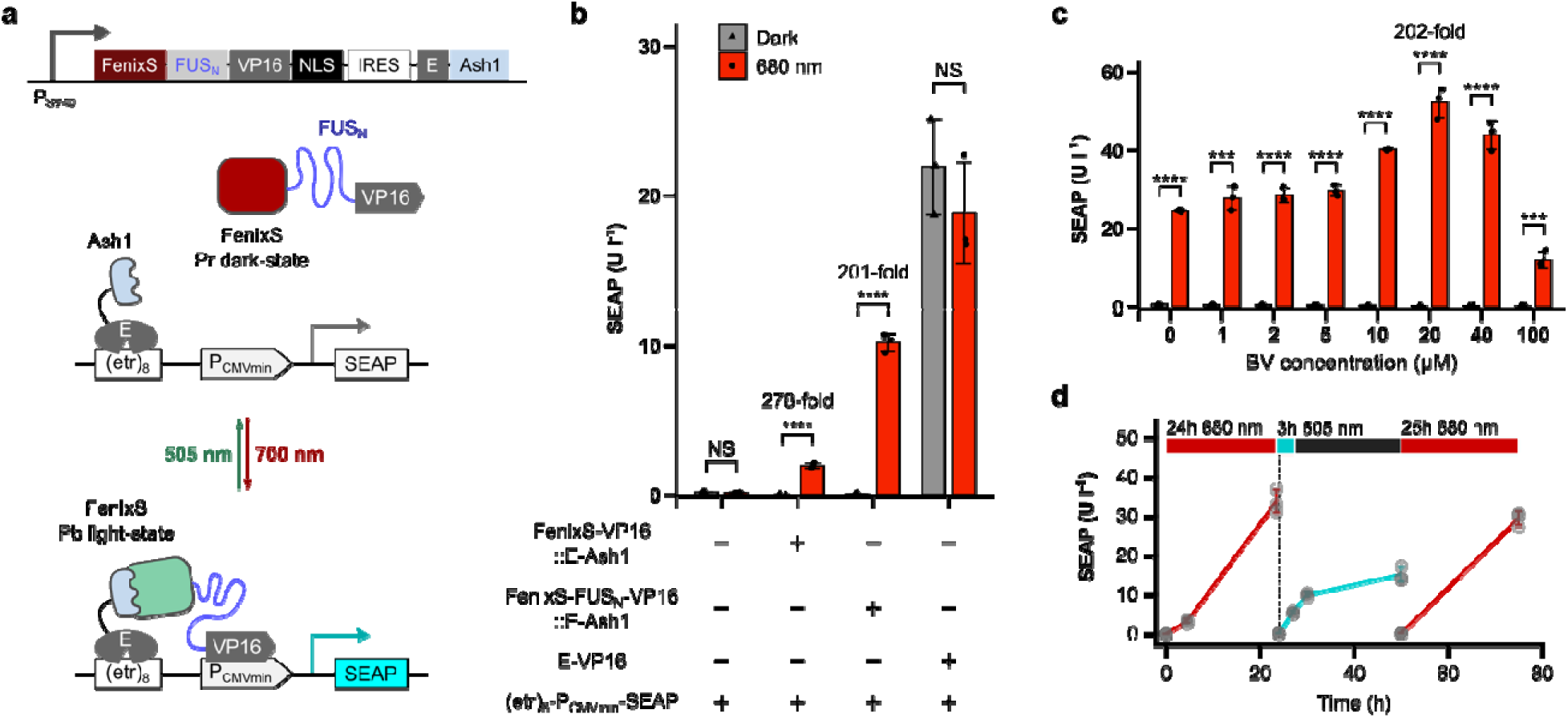
Red light-inducible gene expression using the FenixS-Ash1 system. **a**, Construct design for red light-induced transcriptional activation based on FenixS-Ash1. Upon red light illumination, FenixS and Ash1 associate, recruiting the transcriptional activator to initiate gene expression. **b**, SEAP expression in CHO-K1 cells transiently transfected with different FenixS-Ash1-based constructs and an equimolar amount of the SEAP reporter plasmid. Cells were either kept in the dark or exposed to red light (680 nm, 0.59 mW cm^-2^). FUS_N_ refers to the N-terminus (amino acid 1-214) of the human oncogene FUS, fused to the C-terminus of FenixS. n = 3 biological independent samples, mean ± sd. **c**, SEAP expression in CHO-K1 cells transfected with the FenixS-FUS_N_-VP16-E-Ash1 construct and SEAP reporter plasmid, under varying concentrations of biliverdin (0-100 µM) added to the culture medium. **d**, Reversible control of SEAP expression. Approximately every 24 h, the cell culture medium was replaced with fresh medium supplemented with 20 µM BV and the cells were illuminated as indicated. Cyan light was used to turn off transcription (505 nm, 0.17 mW cm^-2^). **b,c**, Statistical significance was determined using a two-tailed unpaired *t*-test comparing dark versus light conditions: NS (not significant, P > 0.05), *\*P* < 0.05, *\*\*\*P* < 0.001, *\*\*\*\*P* < 0.0001.

Finally, the FenixS-Ash1 system enabled fully reversible control of SEAP expression under alternating red and cyan light cycles (Fig. 4d). These results confirm that the FenixS-Ash1 system functions as designed, enabling wavelength-specific, reversible transgene expression with minimal background activity.

### Control of subcellular localization

Subcellular protein localization is essential for numerous cellular processes. For instance, SOS-recruitment to the plasma membrane is crucial for Ras activation in MAPK pathway^45^; nuclear translocation of NF-κB initiates transcription of target genes in the NF-κB pathway^46^; and translocation of BAX to mitochondria in the BCL-2 family signaling pathway induces cell apoptosis^47^. To examine the performance of the FenixS-Ash1 system in light-controlled protein localization, we performed mitochondrial co-localization experiments. For these experiments, 5 µM biliverdin was added to the culture medium 3 hours prior to imaging to ensure sufficient holo-protein levels.

In these experiments, Ash1 was tagged with mVenus and anchored to the mitochondrial outer membrane via an N-terminal NTOM20 tag, while FenixS was tagged with mCherry and expressed cytosolically (Fig. 5a). Mitochondria-to-cytoplasmic fluorescence ratios (MCRs) were calculated to assess light-inducible protein recruitment. In the dark, mCherry-FenixS was excluded from mitochondria, indicating negligible affinity of Ash1 for the apo and Pr state of FenixS. Upon red light illumination, we observed the localization of FenixS to mitochondria, which was then released upon green light illumination for at least four cycles (Fig. 5b,c and Movie S2). Increased BV concentrations enhanced mitochondrial recruitment of FenixS (Extended Data Fig. 7), and similar localizations patterns were observed with weaker binders Ash0.8 and Ash0.9 (Supplementary Fig. 14). We note that photoconversion of FenixS was achieved using standard fluorescence imaging settings: Pr-to-Pb conversion with 680 nm light (15% laser power, 10 frames, 2.8 s/frames) and Pb-to-Pr conversion using 500 nm light (3% laser power, 15 frames, 2.8 s/frame), which also served to image mVenus.

**Fig. 5:**
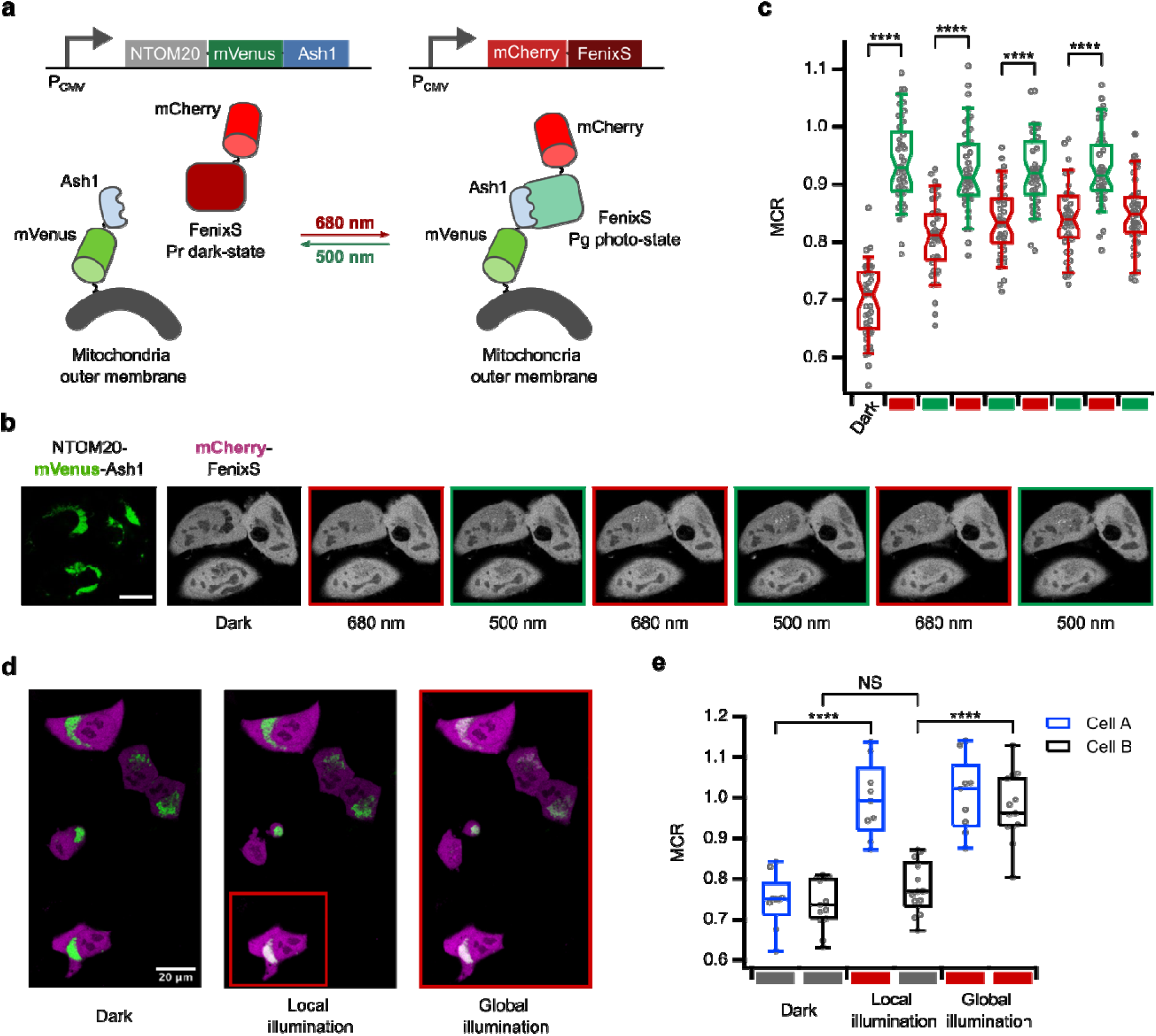
Light-controlled colocalization of FenixS with mitochondria-targeted Ash1 in Hela cells. **a,** Schematic of the construct design and light-controlled recruitment of FenixS to mitochondria-targeted Ash1. **b,** Representative confocal fluorescence images of mCherry-FenixS and NTOM20-mVenus-Ash1 in Hela cells. FenixS was toggled between its dark-and photo-states using alternating green (500 nm) and red (680 nm) light. Scale bar, 20 µm. **c,** Quantification of the mitochondrial to cytoplasmic mCherry fluorescence ratio (MCR) across multiple red-green light cycles (left to right). n = 43 cells from 3 independent experiments. *\*\*\*\*P* < 0.0001 (two-tailed *t*-test, 95% confidence interval). The centre notch of the box indicates the 95% confidence interval of the median; box edges represent the 25^th^ and 75^th^ percentiles; whiskers indicate the 9^th^ and 91^st^ percentiles. **d,** Representative confocal images from a spatially controlled colocalization experiment. Localized 680 nm illumination (red box) induced recruitment of mCherry-FenixS to mitochondria in the targeted region (cell A), without affecting neighboring regions (cell B). Scale bar, 20 µm. **e,** Quantification of MCR for cell A (locally illuminated) and cell B (neighboring cells) under both local and global red-light illumination. n = 9 cells (cell A) and n = 13 cells (cell B) from 3 independent experiments. *\*\*\*\*P* < 0.0001; NS, not significant (two-tailed *t*-test, 95% confidence interval). The centre line of the box indicates the median; box edges represent the 25th and 75th percentiles; whiskers indicate the minimum and maximum values.

A key advantage of optogenetic tools is their ability to control molecular processes with high spatiotemporal precision. To demonstrate spatial control, we selectively illuminated a single cell (Cell A) with red light, resulting in mitochondrial localization of FenixS, while a neighboring cell (Cell B) remained unaffected (Fig. 5d, middle image). Global red light illumination induced mitochondrial localization across all cells in the field of view (Fig. 5d, right image; Fig. 5e).

### Red light-activable intracellular droplet condensation

We next evaluated the applicability of the FenixS-Ash1 pair for red light-controlled condensate formation. Biomolecular condensates are membrane-less compartments formed via liquid-liquid phase separation, concentrating proteins and nucleic acids to spatially and temporally regulate biochemical reactions. These condensates play critical roles in RNA metabolism, stress responses, and transcription regulation^48,49^. Given their significance, several optogenetic tools have been developed to dissect the roles of condensates^50^. The blue light-responsive systems LARIAT^51^, optoDroplet^52^, and Corelets^19^, have demonstrated robust control over condensate formation and dissolution in mammalian cells. However, due to the properties of their blue light-responsive cores, condensates rapidly dissolve after blue light removal – within 2 minutes for Corelets and 10 minutes for LARIAT and optoDroplet. Red/far-red light-responsive system based on *Arabidopsis thaliana* Phytochrome B (PhyB) have also been used to induce condensate formation^53^. However, PhyB-mediated condensation is restricted to temperatures at or below 20°C, limiting its utility in mammalian physiological conditions^53^.

To overcome this limitation, we created R-Corelets by replacing the iLID-SspB pair in Corelets^19^ with the FenixS-Ash1 pair. Specifically, FenixS was fused to FTH1, a 24-mer ferritin assembly, and Ash1 was fused to FUS_N_ domain, an intrinsically disordered region (IDR) of the human oncogene FUS (Fig. 6a). Upon red light illumination, Ash1-FUS_N_ is recruited to the FTH assembly, locally exceeding the saturation concentration of the IDR and triggering phase separation (Fig. 6a). Both components of R-Corelets system were sequentially introduced into HEK293T cells via lentiviral transduction. As anticipated, red light (680 nm) induced condensate formation only in cells expressing both constructs (Fig. 6b). Condensates appeared within ∼10-20 s of illumination and exhibited concentration-dependent kinetic behavior (Fig. 6c). As with Corelets, R-Corelets formed liquid-like condensates that rapidly fused with one another and rounded upon contact (Fig. 6d, Movie S3). Importantly, unlike blue light-responsive condensates based on Cry2-CIB1^51^ or iLID-sspB^19^, which rapidly dissociate upon light removal, the slow thermal relaxation of FenixS-Ash1 sustains condensates post illumination. Condensates remained stable for at least 10 minutes after red light removal, with no observable decrease in intensity or distribution (Fig. 6e). 3D imaging revealed uniform nuclear distribution of condensates (Fig. 6f, Movie S4). These results demonstrate that the FenixS-Ash1 pair enables red light-induced condensate formation, offering a thermally stable and physiologically compatible optogenetic tool for probing phase separation in living cells.

**Fig. 6:**
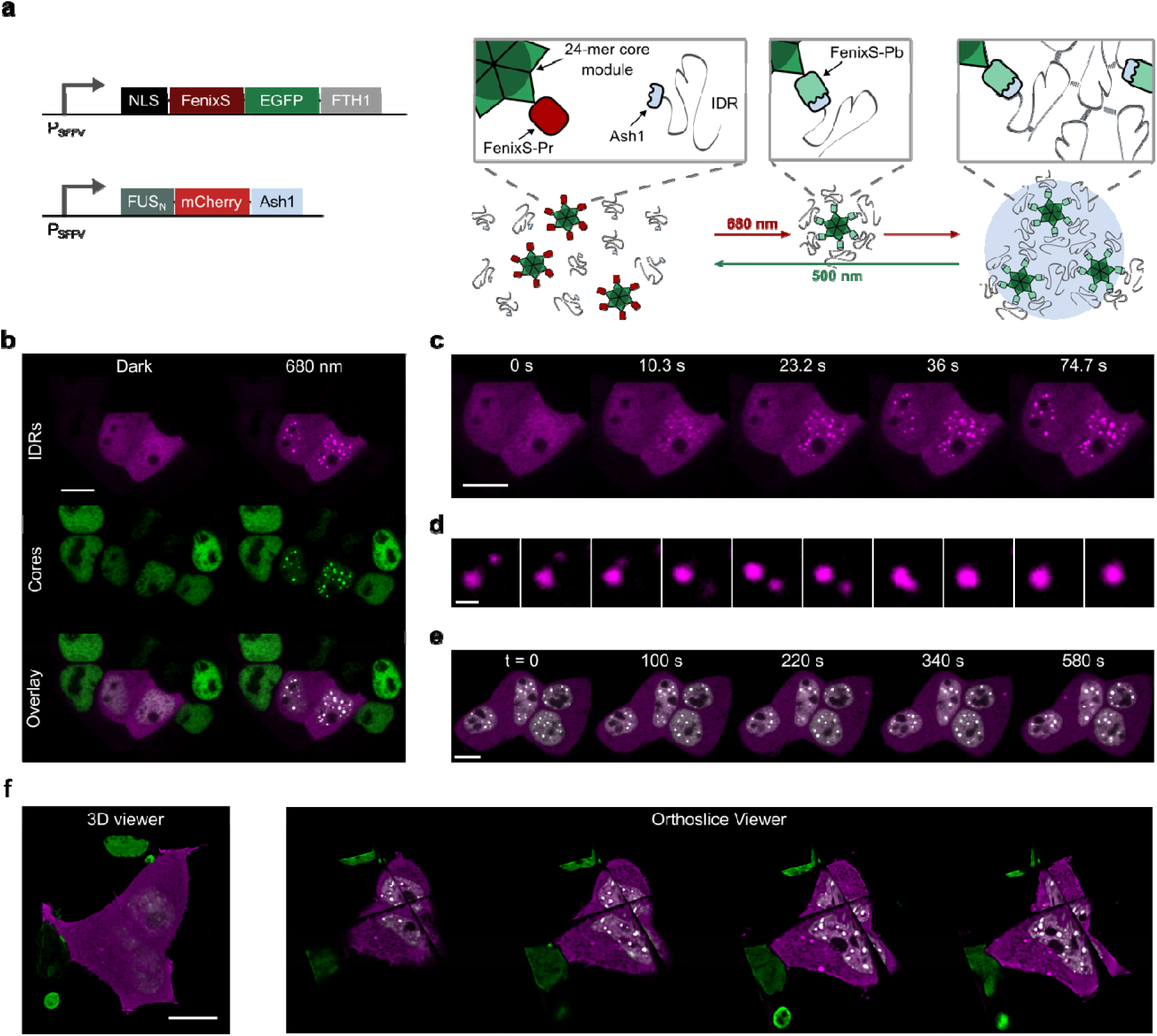
Red light-induced condensate formation using the R-Corelet system. **a**, Schematic of the R-Corelet design, adapted from Bracha et al., 2018^19^. The system comprises two modules: (1) a nuclear-localized, EGFP-tagged ferritin core carrying 24 copies of FenixS, and (2) a mCherry-tagged, photo-state-specific binder Ash1 fused to the intrinsically disordered FUS_N_ domain. Both components were genomically integrated into HEK293T cells via lentiviral transduction. Upon red light activation, FenixS undergoes a conformational change and recruits up to 24 copies of Ash1-FUS_N_ to the core, locally exceeding the saturation concentration of the IDR and triggering phase separation. **b**, Representative confocal fluorescence images of EGFP-tagged core (green) and mCherry-tagged IDRs (magenta) in HEK293T cells expressing both R-Corelet components. Red light (680 nm) induces condensate formation with clear colocalization of the core and FUS_N_ IDR only the cells that expressed both components. Scale bar: 10 µm. **c**, Time-lapse images of condensates formation in the same cell, showing concentration-dependent kinetics. Scale bar: 10 µm. **d**, R-Corelet condensates exhibit liquid-like bahavior, rapidly fuse and rounding upon contact (2.6 s between frames). Scale bar: 1 µm. See also Movie S3 for whole-cell fusion dynamics. **e**, Condensates remains stable for at least 10 minutes following red light removal. Scale bar: 10 µm. **f**, 3D reconstruction of fluorescence images showing uniform nuclear distribution of condensates (left: 3D Viewer; right: Orthoslide Viewer). Scale bar: 10 µm. See also Movie S4 for full 3D visualization.

## Discussion

We have developed a compact, biliverdin-binding optogenetic system based on a cyanobacteriochrome GAF domain for red light-inducible heterodimerization, designated FenixS-Ash1. FenixS was engineered through an intensive protein design and directed evolution campaign. The initial prototype was derived from miRFP670nano by introducing five mutations surrounding D-ring of BV to accommodate the isomerization required for CBCR-GAF domain photoswitchability. Using reversible fluorescence as a high-throughput screening strategy, we iteratively evolved this prototype to yield FenixS – a photoswitchable CBCR-GAF domain with efficient BV binding and robust red/blue photoconversion. This represents the first example of a photoswitchable CBCR-GAF domain engineered through directed evolution. Given the natural diversity of CBCR-GAF domains^20^, we expect them to be highly amenable for engineering. FenixS can be further evolved to enhance BV-binding efficiency or to shift spectral properties by coupling directed evolution with high-throughput screening methods such as flow cytometry.

Several key mutations emerged during evolution. Notably, V95C introduces a cysteine residue near the chromophore. While a blue-absorbing state in a CBCR GAF domain has previously been attributed to the formation for a second thioether bond at the C10 position of the chromophore^33^, the location of Cys95 is novel. If this mutation indeed facilitates C10-adduct formation, it would imply that significant structural rearrangement occurs in the chromophore binding pocket upon isomerization. We note that the large spectral separation between the red and blue states of FenixS makes it a promising candidate for photoacoustic imaging applications, which benefit from high contrast in absorbance between dark and photo states^54^. Another mutation, R98S, decreased fluorescence in the Fenix R18 variant while significantly improving BV-binding efficiency. Further structural studies using X-ray crystallography, molecular dynamics simulations, and mutagenesis will be needed to elucidate the mechanistic role of this mutation.

After developing FenixS, we developed a photo-state-specific binder via phage display. Ash1 binds the photo state of FenixS with a *K*_d_ of 41 nM – surpassing the tightest interaction in the BICYCL series (BICYCL-Green2.4, *K*_d_ = 120 nM) and other BV-binding optogenetic systems such as MagRed (*K*_d_ = 310 nM) and NanoRED (*K*_d_ = 470 nM). This affinity is comparable to state-of-the-art blue light-responsive systems, LOVTRAP (AsLOV2-Zdk1, *K*_d_ = 26 nM). With an estimated *K*_d_ > 50 µM for the OFF state (dark state), the FenixS-Ash1 pair exhibits a > 1,200-fold change in affinity between ON and OFF states – the largest dynamic range reported for any heterodimerization system to date. This high affinity and dynamic range make FenixS-Ash1 suitable for a wide range of applications with varying concentration threshold requirements for their activity, eliminating the need for multiple tools with different affinity ranges (e.g. iLID nano and iLID micro systems^7,17^).

We hypothesize that the exceptional dynamic range of FenixS arises from substantial structural rearrangement upon photoconversion, potentially driven by C10-adduct formation. In the photo state, FenixS binds tightly to Ash, while in the dark state, the absence of key structural features prevents association, resulting in ultra-low background activity. Further studies will provide deeper insight into the structural basis of the observed light-dependent binding specificity.

The versatility of FenixS-Ash1 was demonstrated across multiple experimental platforms. In co-localization experiments, FenixS-Ash1 enables light-inducible recruitment of proteins to agarose-bead surfaces and to mitochondrial outer membranes. Using FenixS-Ash1, we developed R-Corelets based on the design of Corelets, thus enabling red light-induced condensate formation. In transcriptional activation assays, FenixS-Ash1-based optogenetic tools exhibit ∼200-fold induction of gene expression upon red light activation. Notably, 47% of the maximal activation was achieved without exogenous BV, highlighting a major advance in engineering BV-binding CBCR-based optogenetic tools. Although miRFP670nano functions with endogenous BV, subsequent studies indicated that miRFP670nano is not fully chromophorylated^25^, and like other phytochrome-based fluorescence proteins, brightness is enhanced by BV supplementation or co-expression of a BV-synthesizing enzyme^44^. Whether BV supplementation is beneficial for FenixS-Ash1 performance is expected to depend on the organism and experimental context and should be empirically determined for each application. Importantly, Ash1 does not bind apo-FenixS, ensuring that non-chromophorylated protein – commonly present in heterologous systems – does not interfere with system performance. Finally, FenixS exhibits robust photo-reversibility with repeated association/dissociation cycles using 700-nm and 505-nm illumination, while slow thermal reversion enables sustain activation after red light is removed.

## Supporting information

Description of Addtional Movie Files

## Acknowledgements

The authors are grateful to N. Elsakrmy and H. Cui (University of Toronto) for providing access to equipment and technical assistance, W. Walker and J. Jonkman (The Advanced Optical Microscopy Facility, UHN) for support with imaging experiments, K. Tang and M. T. Zurbriggen (University of Düsseldorf) for the kind gift of transcriptional plasmids, V. Vologodskaya for technical assistance, and J. Lee and H.O. Lee (University of Toronto) for helpful discussions.

## Author contributions

G.N.T.L., G.A.W., and M.U. conceived the project. G.N.T.L. and L.M.T.P designed and carried out the directed evolution of FenixS. G.N.T.L designed, carried out and analyzed all experiments in vitro and in mammalian cells. B.X. assisted with the purification of FenixS. G.A.W. and M.U. supervised the project. G.N.T.L. and G.A.W. wrote the manuscript and prepared the figures. All authors edited and approved the final manuscript.

## Methods

### General Methods and Materials

Q5 high-fidelity DNA polymerase (New England Biolabs) was used for routine polymerase chain reaction (PCR) amplification, and Taq DNA polymerase (New England Biolabs) was used for error-prone PCR. Primers for DNA amplification and site-directed mutagenesis were synthesized by Integrated DNA Technologies (IDT). Restriction endonucleases (New England Biolabs) and T4 DNA ligase (New England Biolabs) were used for plasmid construction. Products of PCR and restriction digests were purified using agarose gel electrophoresis and the GeneJET gel extraction kits (Thermo Fisher Scientific) or Monarch Spin PCR & DNA cleanup kit (New England Biolabs). Lentiviral plasmids were cloned using NEBuilder HiFi DNA assembly master mix (New England Biolabs) in a standard reaction mixture comprising 20 ng of each 2-3 PCR-amplified inserts and 40 ng backbone in a 4 µl reaction. Bacterial amplified plasmids were isolated via plasmid DNA miniprep kits (Thermo Fisher Scientific) for non-mammalian cell works, plasmid DNA midiprep (Thermo Fisher Scientific) for mammalian cell work, and QIAGEN plasmid midi kits for lentiviral work. DNA sequences were analyzed by DNA sequence service of TCAG and whole plasmid sequencing by Plasmidsaurus. The fluorescence spectra and intensity were recorded on a BioTek Synergy H1 microplate reader or CLARIOstar Plus microplate reader (BMG LABTECH). The UV-Vis absorbance spectra were obtained on SPECTROstar Nano microplate reader (BMG LABTECH) using a 10 mm quartz cuvette (Hellma Analytics). All plasmids in this study are listed in the Supplementary Table 1.

### Directed evolution of FenixS

The gene encoding miRFP670nano^24^ (a gift from Vladislav Verkhusha; Addgene plasmid #127427) and miRFP670nano3^25^ (Addgene plasmid #184663) with an N-terminal 6x His tag was cloned into the pBAD/HisB vector^55^ (a kind gift from Robert E. Campbell). This plasmid also contains the gene encoding heme oxygenase 1 (HO1), which catalyzes the conversion of heme to biliverdin in *E. coli*. Protein variants were expressed in *E. coli* strain 10-beta (New England Biolabs) grown in LB medium supplemented with 100 µg mL^-1^ ampicillin and 0.02% L-arabinose for 18 hours at 37°C. Proteins were extracted using B-PER bacterial protein extraction reagent (Thermo Fisher Scientific), followed by six freeze-and-thaw cycles. Fluorescence brightness and photoswitching assays were performed under red (660 nm) and green (530 nm) illumination. During directed evolution, MnCl_2_ concentration was adjusted to achieve an average of 1-2 mutations in the whole gene. Primary screening was done on the agar plates, evaluating ∼ 25,000 colonies per round. Fluorescence images were captured using a SYNGENE G:BOX (excitation: 640 – 660 nm; emission: 697 – 717 nm). OFF-state images were taken after 660 nm irradiation (5 minutes, 12 mW cm^-2^), and ON-state images were taken after 530 nm irradiation (5 minutes, 1 mW cm^-2^). The fluorescence images of the ON and OFF states were overlaid and analyzed using ImageJ. A total of 96 colonies exhibiting desirable traits were selected for protein extraction and fluorescence measurement. After 18 rounds of evolution, the introduction of the R98S mutation ultimately yield final variant, FenixS.

Fluorescence measurements were performed using a BioTek Synergy H1 microplate reader using a Greiner 96-well flat-bottom microplate. Cell lysates in HEPES buffer were added to 96-well plate, and fluorescence was measured at Ex/Em = 647/670 nm. OFF-state fluorescence was recorded after red light irradiation (660 nm, 10 minutes, 12 mW cm^-2^), followed by green light irradiation (530 nm, 10 minutes, 1 mW cm^-2^) for ON-state. At least two photoswitching cycles were performed to assess fatigue resistant, selecting variants that maintained ON-state fluorescence across cycles.

### Protein expression and purification

To produce holo-FenixS via *in vivo* assembly, *E. coli* 10-beta cells were transformed with the pBAD-FenixS-HO1 plasmid. A single colony was inoculated in 5 mL LB (100 μg mL^-1^ ampicillin) and grown overnight at 37 °C. The saturated culture (1 mL) was used to inoculate 1 L of LB (100 μg mL^−1^ ampicillin) and cells were grown at 37 °C until reaching and OD_600_ of 0.6. Protein expression was induced by adding 200 mg L-(+)-arabinose, followed by incubation for 5 hours at 37 °C with shaking at 180 rpm. Cells were harvested by centrifugation at 4 °C, 4,000 rpm for 1 h. Cells resuspended in lysis buffer (10 mM HEPES, 150 mM NaCl, 1 mM DTT, 5 mM benzamidine, pH 7.4) were sonicated and centrifuged to remove cell debris. Lysates were filtered with a 0.45 µm filter and loaded onto a lysis buffer pre-equilibrated Ni-NTA column. The column was washed extensively with wash buffer (10 mM HEPES, 150 mM NaCl, 5 mM benzamidine, pH 7.4), and holo-FenixS was eluted using elution buffer (10 mM HEPES, 150 mM NaCl, 400 mM imidazole, pH 7.4). The eluted protein was dialyzed twice against 1L dialysis buffer (10 mM HEPES, 150 mM NaCl, 1 mM EDTA, pH 7.4). Holo-protein was further purified by size exclusion chromatography using a Superdex 75 10/300 GL increase column (GE Healthcare) and confirmed via electrospray ionization mass spectrometry (ESI-MS).

For the preparation of apo-FenixS, apo-AM1_C0023g2 S334G^12^, and apo-miRFP670nano, the corresponding genes were cloned into pET24b expression vectors containing a N-terminal 6x His tag and were expressed in *E. Coli* BL21 (DE3) (New England Biolabs). A single colony was inoculated in 5 mL LB culture (50 μg mL^-1^ kanamycin) and grown overnight at 37 °C, shaking at 180 rpm. Saturated culture (1 mL) was used to inoculate 1 L of LB (50 μg mL^−1^ kanamycin), and cells were grown at 37 °C until OD_600_ reached 0.6. Protein expression was induced with 1 mM IPTG, followed by overnight shaking at 18 °C. The next day, cells were harvested and purified using the protocol above, with the exception that all buffers were supplemented with 1 mM DTT to maintain reducing conditions.

For phage display, a C-terminal AviTag sequence was introduced into pBAD-FenixS-HO1 plasmid (see above). The protein expression and purification were performed using the same procedures as described above, up to the dialysis steps. To ensure complete chromophore incorporation, an excess amount of biliverdin (Frontier Scientific), dissolved in DMSO, was added to the protein solution. The mixture was incubated at room temperature for an hour, followed by overnight incubation at 4°C. Excess BV was removed by dialysis against dialysis buffer. Biotinylation of AviTag-holo-FenixS was performed according to a published protocol^56^. The reaction mixture containing 100 µM AviTag-holo-FenixS in 5 mM Mg(OAc)_2_, 2 mM ATP, 1 µM BirA enzyme, and 150 µM D-Biotin. The mixture was incubated for 1 hour at room temperature, followed by overnight incubation at 4 °C. Biotinylated holo-FenixS was further purified by size exclusion chromatography as described above and its mass was confirmed via electrospray ionization mass spectrometry (ESI-MS). To prepare biotinylated apo-FenixS, the gene of FenixS was cloned in pET24b vector containing a C-terminal AviTag and an N-terminal 6x His tag, and expressed in *E. coli* BL21(DE3). Expression and purification were carried out using the same protocol as for apo-FenixS. Biotinylation of AviTag apo-FenixS was performed using the same procedures as described for the holo form. For long-term storage, biotinylated protein in storage buffer (10 mM HEPES, 150 mM NaCl, 1 mM EDTA, 10% glycerol, pH 7.2) was flash frozen using liquid nitrogen and stored at -80°C.

### Biliverdin binding assay

Purified apoprotein (7.5 µM) was mixed with 5 µM biliverdin (BV; 100x stock in DMSO) in HEPES buffer containing 1 mM DTT at 25°C. Absorbance spectra were recorded immediately after mixing (within 5 seconds) and at subsequent time points until the absorbance reached a steady state. The absorbance at the Q-band peak was plotted as a function of time and fitted to a single-exponential association model to determine the BV-binding rate: Abs(t) = c + A*(1-exp(-(ln(2)/t_1/2_)*t)).

### Size exclusion chromatography

Size-exclusion liquid chromatography of FenixS and binders (Ash1, Ash0.8, Ash0.9) were performed using Superdex 75 10/300 GL size exclusion column (GE Healthcare, 0.4 ml min^−1^ flow rate). The column was equilibrated with HEPES buffer (10 mM HEPES, 150 mM NaCl, 1 mM EDTA, pH 7.4). All proteins were injected at a concentration of 50 µM and a volume of 0.25 mL. The column was calibrated with Bio-Rad gel filtration standards (10 mg ml^-1^ Thyroglobulin (bovine), 10 mg ml^-1^ g-globulin (bovine), 10 mg ml^-1^ Ovalbumin (chicken), 5 mg ml^-1^ Myoglobin (horse), 1 mg ml^-1^ Vitamin B12).

### UV-Vis spectroscopy

For native absorbance spectra, 5 µM FenixS in HEPES buffer (10 mM HEPES, 150 mM NaCl, 1 mM EDTA, pH 7.4) was used. Samples were either kept in the dark to obtain the Pr state or irradiated with 700 nm light to generate Pb state. For denatured absorbance spectra, 50 µM FenixS in HEPES buffer was either kept in the dark or irradiated with specific wavelengths (455 nm, 505 nm, 530 nm, or 700 nm). The samples were then diluted to 5 µM in denaturation buffer (8 M Gdn.HCl, 200 mM NaOAc, pH 4.4) prior to measurement.

### Phage display-based selection

For the naïve selection, a GA domain naïve library displayed on the M13 phage pVIII coat protein (as previously described^36^) was used to identify binders specific to the Pb state of FenixS, following an optimized protocol published earlier^34^. To generate the Pr and Pg state of FenixS, selection plates were illuminated with green (530 nm) and red (660 nm) light, respectively. After three rounds of selection, 96 individual colonies were screened using phage-based single clone ELISA (detailed previously published protocol^34^), leading to the identification of the initial hit, Ash0.1.

For affinity maturation, a biased library was constructed based on doped oligonucleotides specific for lead clones Ash0.1 and displayed on the M13 phage pIII coat protein. The following oligonucleotide was used for mutagenesis:

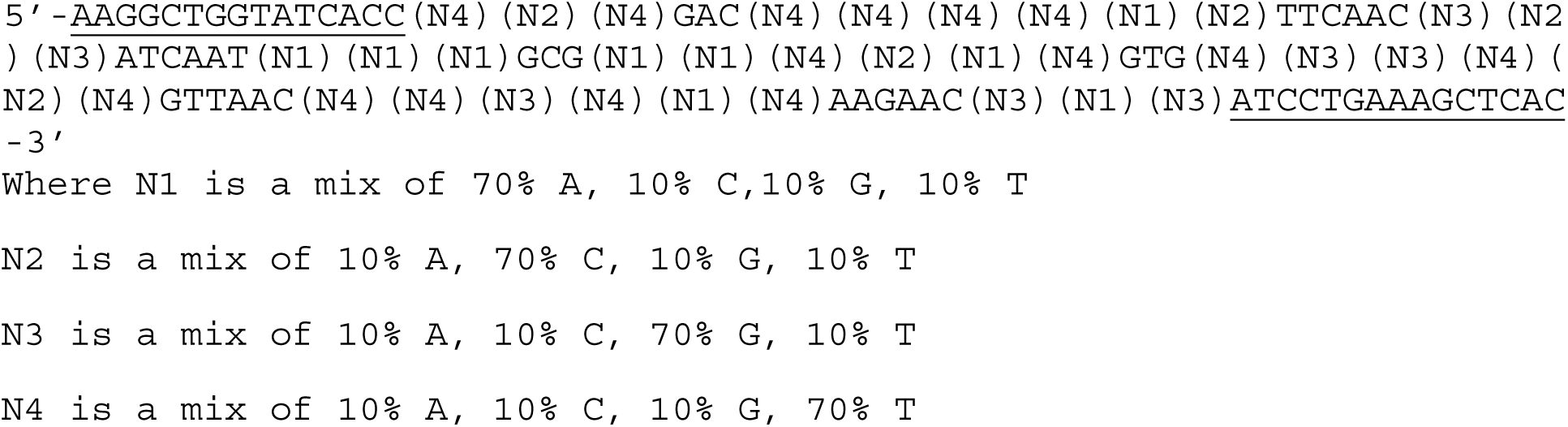

The biased library was constructed following previously published protocols, resulting in an estimated diversity of approximately 10^9^ different clones^57^. Affinity maturation selection was performed using the same general protocol as the naïve selection, with specific modifications to the first negative selection steps. In round one, the first negative selection was carried out by incubating the phage library with soluble apo-FenixS at two concentrations: 50 nM (condition #1) and 250 nM (condition #2). For condition #1, the concentration was increased to 100 nM in round two and 250 nM in round three. For condition #2, the concentration was increased to 500 nM in round two and 1250 nM in round three. These steps were designed to eliminate clones that bind non-specifically to the apo form of FenixS. The second negative selection step against Pr state and the subsequent positive selection for the Pb state were performed identically to naïve selection protocol. After three rounds of selection, 96 individual colonies were screened using phage-based single clone ELISA to identify improved binders.

### ELISA of purified binders

Selected binders containing an N-terminal FLAG-tag were amplified from pIII phagemids via PCR and cloned into the pET24b expression vector containing a C-terminal 6x His tag using XhoI and HindIII restriction sites. The resulting plasmids were used to transformed *E. coli* BL21(DE3), and protein expression and purification were performed using the same protocol as for apo-FenixS.

Titration ELISA of purified binders was carried out as previously described^34^. Two MaxiSorp 96-well plates were coated with 2 µg ml^-1^ streptavidin in PBS (50 µl per well) and blocked with 100 µl PB buffer (PBS with 2 mg ml^-1^ BSA). After washing with PT buffer (PBS with 0.05% Tween-20), each plate was coated with 400 nM (50 µl) of holo-FenixS (rows A-D), apo-FenixS (rows E-F), or PBT buffer as negative control (row G-H). Plates were incubated for 1 hour at room temperature. After removing unbound proteins, 50 µL of serial diluted binder (ranging from 50 µM to 1 nM) was added to each well. One plate (Pr state) was placed under green light (530 nm), and the other (Pb state) was placed under red light (660 nm). After 1 hour incubation at room temperature under their respective light conditions, plates were washed with PT buffer. Then, 50 µL of anti-FLAG-HRP antibody (Antibody Design Labs; diluted 1:10,000 in PBT buffer) was added to each well and incubated for 1 hour under their respective light conditions. Following antibody incubation, plates were washed with PT buffer, and 50 µl of TMB substrate (Thermo Fisher Scientific) was added to each well. After 5 minutes of color development, the reaction was stopped with 50 µl of 1 M phosphoric acid. Absorbance at 450 nm was measured. ELISA signals as a function of binder concentration were fitted to a dose-response equation (f(x) = base + (max-base)/(1+*K*_d_/x)) to determine the apparent dissociation constant *K*_d_.

### Circular dichroism (CD)

Circular dichroism experiments were carried out using a Jasco J-710 circular dichroism spectropolarimeter using a 1 mm quartz cuvette (Hellma Analytics). All protein binders were prepared at approximately 50 µM in 10 mM sodium phosphate buffer (pH 7.2) and measurements were taken at 20°C. Accurate concentration was determined based on absorbance at 280 nm directly with sample in CD cuvette using Lambda 1050 UV/Vis Spectrometer.

### Bead imaging

To the pET24b-FLAG-Ash1 plasmid, the N-terminal FLAG-tag was replaced with mCherry using XhoI and BamHI restriction sites. mCherry-Ash1 fusion protein were expressed and purified with a C-terminal poly His (6x) tag. Streptavidin-coated agarose beads (Sigma) were equilibrated by washing three times with HEPES buffer (10 mM HEPES, 150 mM NaCl, 1 mM EDTA, pH 7.4). To functionalize the resin beads, 125 µL of either biotinylated holo-FenixS or biotinylated IgG (Rockland) was incubated with 50 µL of bead slurry for 1 hour in the dark. Excess unbound protein was removed by washing the beads three time with HEPES buffer. One microliter of FenixS-coated beads and 1 µL of IgG-coated beads were mixed with 50 µL of 500 nM mCherry-Ash1 in HEPES buffer. The mixtures were incubated for 1 hour in the dark and then transferred to a 0.17-mm Greiner 96-well flat-bottom microplate for imaging. Bead imaging was performed using a Leica Mica microscope equipped with a built-in LED light source (470 nm, 170 mW; 555 nm, 170 mW; 625 nm, 170 mW) and a 20× air objective lens. mCherry fluorescence was captured using 555 nm excitation (10.4% intensity, 250 ms exposure). Pr-to-Pb photoconversion of FenixS was induced using 625 nm illumination (32.9% intensity) while Pb-to-Pr photoconversion was induced using 470 nm illumination (67.8% intensity). Image analysis was conducted in ImageJ using the Time Series Analyzer V3 plugin. Background subtraction was applied using the sliding paraboloid method to generate the final movie.

### Gene expression in CHO cells

The genes encoding FenixS and Ash1 were subcloned into plasmids pKT214 and pKT753 (kind gift from M. Zurbriggen) using T4 ligation to generate the desired constructs (see **Supplementary Table 1** for details). CHO-K1 cells (kind gift from Trieu Le) were seeded at a density of approximately 70,000 cells per well in 24-well plates containing F12 Nutrient Mixture media (Ham) (Gibco) supplemented with 100 U ml-1 penicillin, 100 U ml-1 streptomycin, and 10% FBS (Invitrogen, Canadian Origin). The next day, cells were transfected with 750 ng DNA mix (equimolar plasmid amounts, w/w) using polyethyleneimine (PEI 25,000; Polysciences). The medium was replaced 4 – 5 hours post-transfection, and cells were incubated at 37°C (5% CO_2_). At 23 h post-transfection, the medium was replaced with pre-warmed medium supplemented with 40 µM biliverdin (Frontier Scientific), unless otherwise specified. At 24 h post-transfection, cells were either kept in the dark or under 680 nm red light (Thorlab LED, 0.59 mW cm^-2^). At 48 h post-transfection, the cell culture medium was collected for SEAP reporter quantification following previously published protocol^61^.

For kinetic experiment, 400,000 CHO-K1 cells were seeded per well in 6-well plates. The cells were transfected with 3 µg DNA mix (equimolar plasmid amount, w/w). At indicated time points, 100 µL of cell culture medium was collect and stored at – 20 °C for SEAP quantification. In reversibility of gene expression experiment, 400,000 CHO-K1 cells were seeded in 6-well plates and transfected as described above. At 24 h post-transfection, the cells were illuminated with 680 nm light (Thorlab LED, 0.59 mW cm^-2^) for 24 h. At 48 hours, the medium was replaced with fresh medium containing 20 µM BV. Thirty minutes prior to medium exchange, the cells were kept under 505 nm light (Thorlab LED, 0.17 mW cm^-2^) for a total of 3 hours, followed by 23.5 h in the dark. At 74 h post-transfection, the medium was again replaced with fresh medium supplemented with 20 µM BV, and the cells were re-illuminated with 680 nm light for the remainder of the experiment. At each indicated time point, 100 µL of cell culture medium was collect and stored at – 20 °C for SEAP quantification.

### Light-induced mitochondrial protein co-localization in Hela cells

The gene encoding FenixS was subcloned into pTRIEX vector^6^ (a gift from K. Hahn, Addgene plasmid 81041) that contain N-terminal mCherry using BamHI and HindIII restriction sites. Binders were subcloned into the same vector containing an N-terminal mVenus tag that contains an NTOM20 mitochondrial localization tag on its N-terminus using BamHI and XhoI restriction site. Hela cells (a gift from H.O. Lee) were cultured in DMEM (Sigma-Aldrich) supplemented with 10% FBS (Invitrogen, Canadian Origin), 100 U ml^-1^ streptomycin, and 100 U ml^-1^ ampicillin.

Cells were seeded into a two-chamber imaging plate (Labtek). The next day, 1.6 µg DNA mix was transfected in equimolar plasmid amounts (w/w) using polyethyleneimine 25000 (Polysciences). The medium was exchanged 4 h after transfection and cells were incubated at 37°C (5% CO_2_). Approximately 3 h prior to imaging, the medium was replaced with pre-warmed medium additionally supplemented with biliverdin (Frontier Scientific) at a concentration of 5 µM if not indicated otherwise. The medium was replaced with pre-warmed imaging media (DMEM with HEPES buffer, no phenol red) immediately before imaging. Imaging was carried out 48 – 72 h after transfection.

Imaging was carried out using a Leica Stellaris 5 confocal microscope with an HC PL Apo x63/1.4 oil immersion objective equipped with a CO_2_ microscope stage incubator under 5% CO_2_ and 37 °C. Fluorescence proteins were excited and FenixS was photoswitched using white light laser (485 – 790 nm). mVenus was excited at 500 nm (3.5%, 9.5 µW µm^-2^) and emission was monitored at 510 – 600 nm, while mCherry was excited at 560 nm (2.5%, 18.8 µW µm^-2^) and emission was monitored at 585 – 680 nm. FenixS photoconversion was induced using 680 nm light 10 frames (15%, 147 µW µm^-2^) and emission was monitored at 695 – 780 nm.

### Droplet condensation control in stable HEK293T cell

#### Construction of stable cell line

The gene encoding FenixS and Ash1 was subcloned into pHR-SFFV by replacing iLID-SspB system in Corelets (kind gift from Clifford Brangwynne; Addgene plasmid #122147, #122148). R-Corelets construct-containing Lentiviruses were produced by cotransfecting HEK293T cells plated on a 10 cm culture dish with the desired DNA constructs (2 µg), psPAX2 (2 µg) (kind gift from H. Cui), and pCMV-VSV-G^58^ (1 µg) (Addgene plasmid #8454) using PolyJet transfection reagent (FroggaBio, kind gift from H. Cui) following manufacture instructions. Viral supernatants were collected and filtered from cell debris using 0.45 µm filter (FroggaBio) within 3 days following transfection. Aliquots of viral supernatant were stored at -80°C for long term uses. To generate stable cell line expressing R-Corelets, HEK293T cells were sequentially transduced with FS1 and FS2. For each transduction step, HEK293T cells were transduced while at 50% confluency on 6-well plates by adding 1 mL of the harvested Lentiviruses and 10 µg/ml polybrene (Millipore Sigma, kind gift from H. Cui). 3 days following transduction, the cells were washed twice with PBS and passaged normally.

#### Light-induced condensate formation

Imaging was performed using a Leica Stellaris 5 confocal microscope equipped with an HC PL Apo x63/1.4 oil immersion objective and a CO_2_ microscope stage incubator maintained at 5% CO_2_ and 37 °C. EGFP, mCherry, and FenixS were excited using a tunable white light laser (485 – 790 nm). mVenus was excited at 495 nm (2 %, 5.4 µW µm^-2^), with emission was collected between 505 – 600 nm. mCherry was excited at 560 nm (3 %, 22.5 µW µm^-2^), and emission was collected between 585 – 680 nm. FenixS photoconversion was induced using 680 nm light 30 frames (15%, 147 µW µm^-2^).

For 3D imaging of cells containing condensates, dual-channel acquisition (495 nm and 560 nm excitation) was performed with optical sectioning at 0.8 µm (pinhole size: 82.3 µm). Z-stacks were acquired with 47 slices at 0.26 µm intervals. Image stacks were processed using the DeconvolutionLab2 plugin in Fiji^59^, with the following setting: synthetic PSF (Gaussian, dimensions 3.0 3.0 3.0; size 200 x 100 x 100; intensity 255.0), and the Richardson-Lucy Total Variation (RLTV) algorithm (interations:10; regularization: 0.01). 3D reconstructions and orthogonal slice views were generated using the 3D Viewer plugin in Fiji^60^.

---

**Extended Data Fig. 1:**
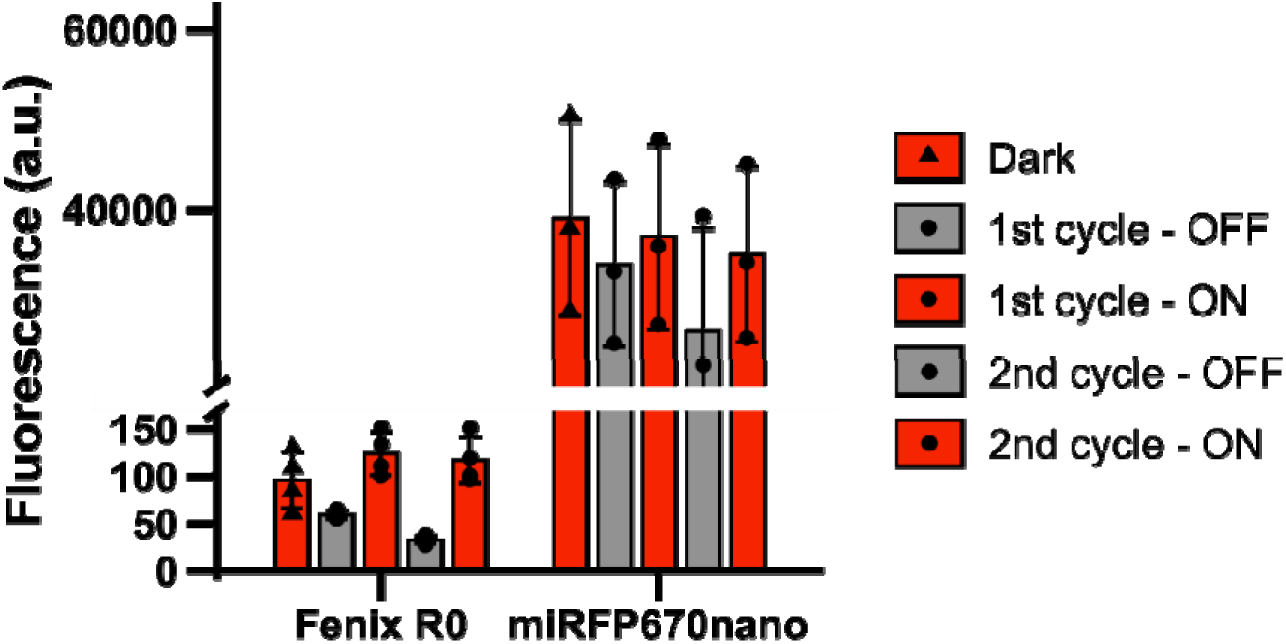
Comparison of photoswitchability of Fenix R0 and miRFP670nano. Fluorescence and photoswitchability of Fenix R0 and miRFP670nano (crude protein extracts from *E. coli*). The OFF-state was produced with 660 nm illumination, the ON-state was produced with 530 nm illumination. Fluorescence of both states was detected at 670 nm with excitation at 647 nm. n = 3 technical replicates, mean ± sd.

**Extended Data Fig. 2:**
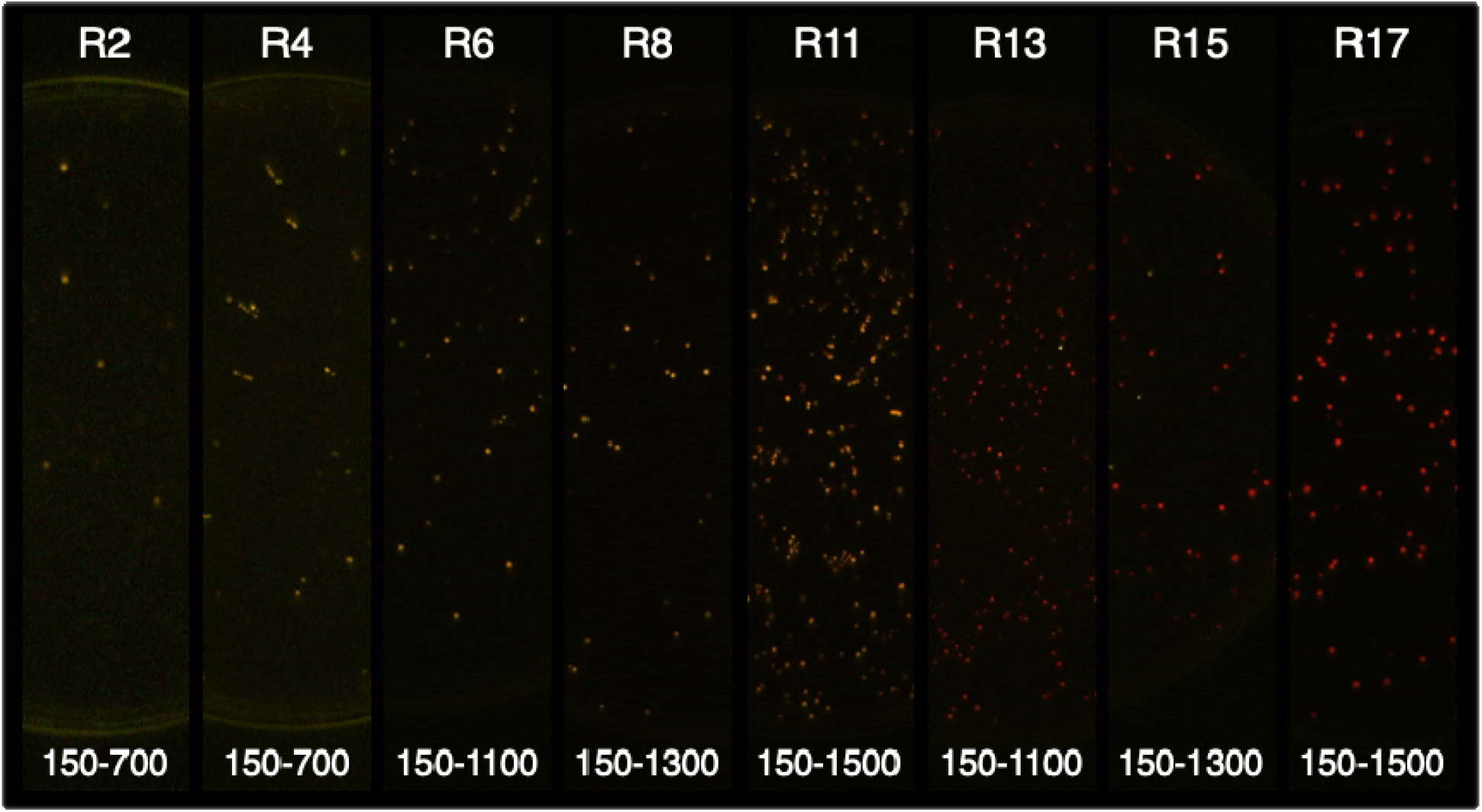
Processed fluorescence images of agar plates during directed evolution. Overlay images of on-state (red-channel) and off-state (green-channel) fluorescence from bacterial colonies throughout the directed evolution process. Progressive increases in brightness and greater difference between on- and off-state fluorescence (redness) indicate enhanced Fenix fluorescence, biliverdin binding efficiency, and photoswitchability. The numbers at the bottom indicate the minimum and maximum brightness values set in ImageJ for image processing.

**Extended Data Fig. 3:**
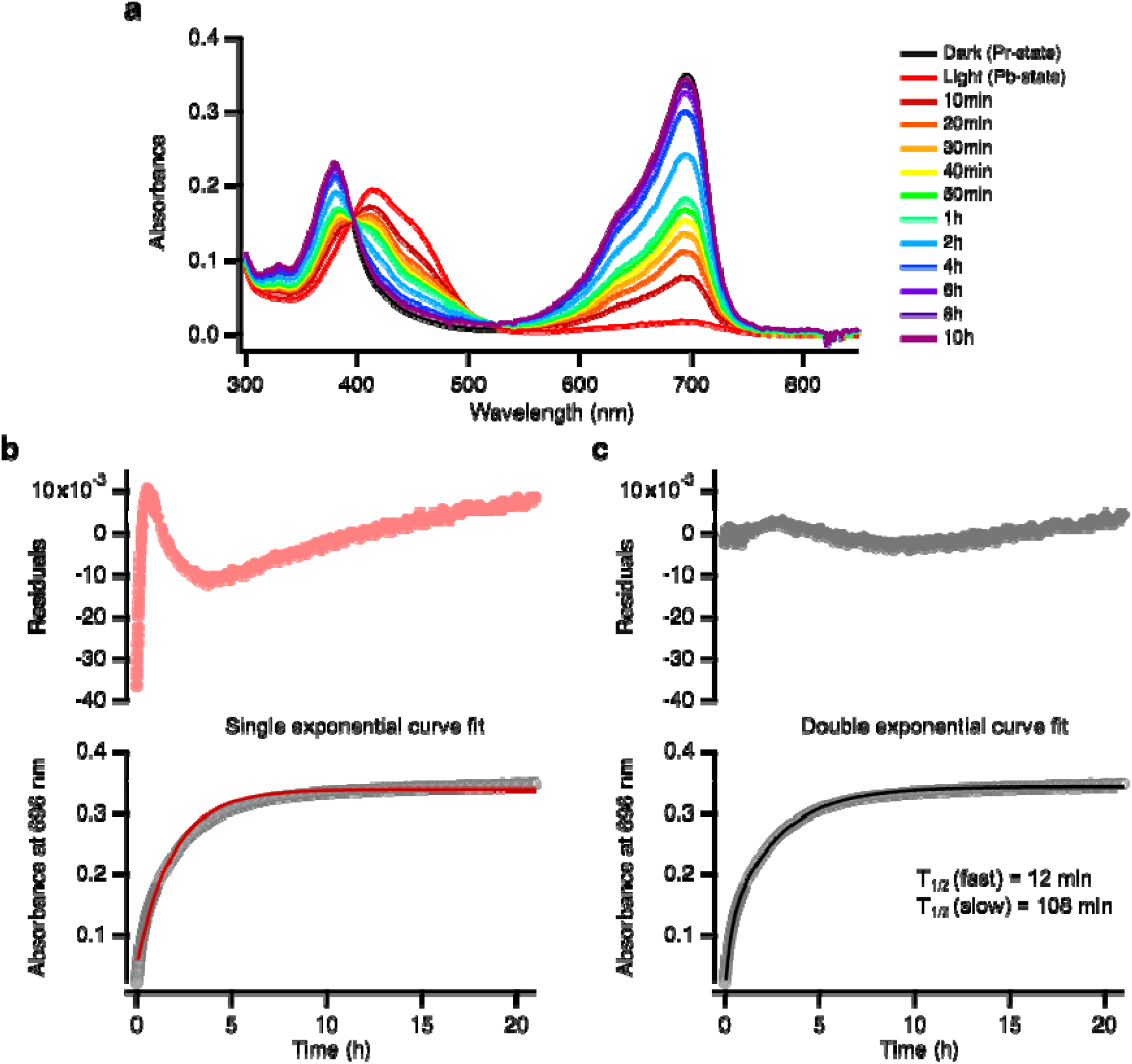
Thermal relaxation of FenixS. **a,** Photoswitching followed by thermal relaxation of FenixS (5 µM) at 37°C. **b,c,** Bottom: absorbance at 696 nm of FenixS as a function of time at 37°C. (b) Single exponential fit: Abs = f(t) = c + A*(1-exp(-(ln(2)/t_1/2_)*t)). (c) Double exponential fit : Abs = f(t) = c + A(fast)*(1-exp(-(ln(2)/t_1/2_(fast))*t)) + A(slow)*(1-exp(-(ln(2)/t_1/2_(slow))*t)). Top panels show residuals from each fit calculated as the difference between fitting function model and raw data. The single exponential fit yields substantially larger residuals, indicating a poor fit compared to the double exponential model.

**Extended Data Fig. 4:**
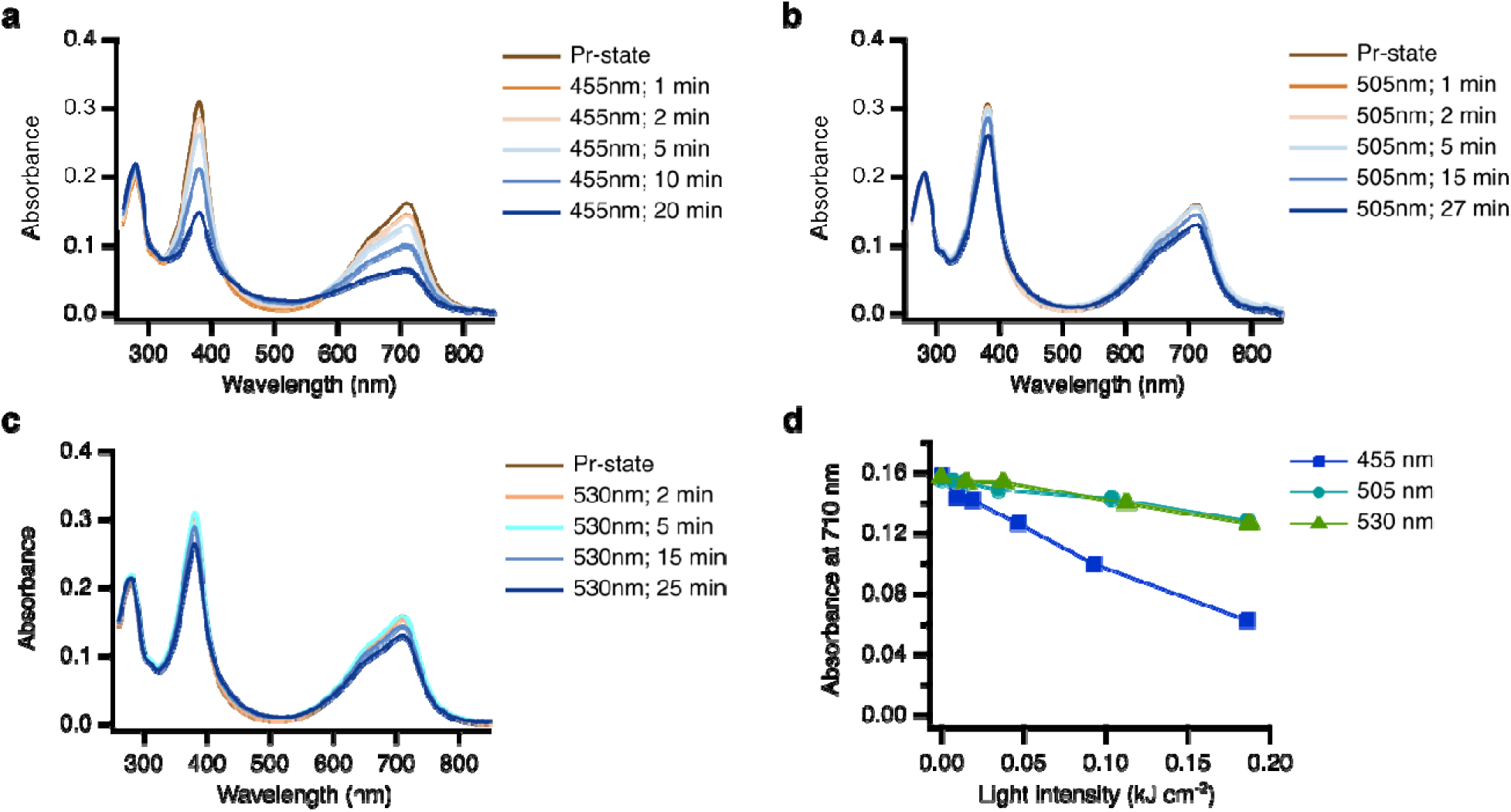
Light-induced photobleaching of the FenixS dark-state. **a-c**. Denatured absorbance spectra of FenixS in the dark state following illumination with (**a**) 455 nm (155 mW cm^-2^), (**b**) 505 nm (115 mW cm^-2^), and (**c**) 530 nm (125 mW cm^-2^) light. A decrease in both the Soret and Q-bands of biliverdin in the denatured spectra indicates photobleaching of the chromophore. **d**. Quantification of photobleaching, monitored by absorbance at 710 nm in denaturing buffer, as a function of light intensity at different wavelengths. Irradiation at 455 nm caused significant photobleaching, whereas this effect was substantially reduced with 505 nm or 530 nm light.

**Extended Data Fig. 5:**
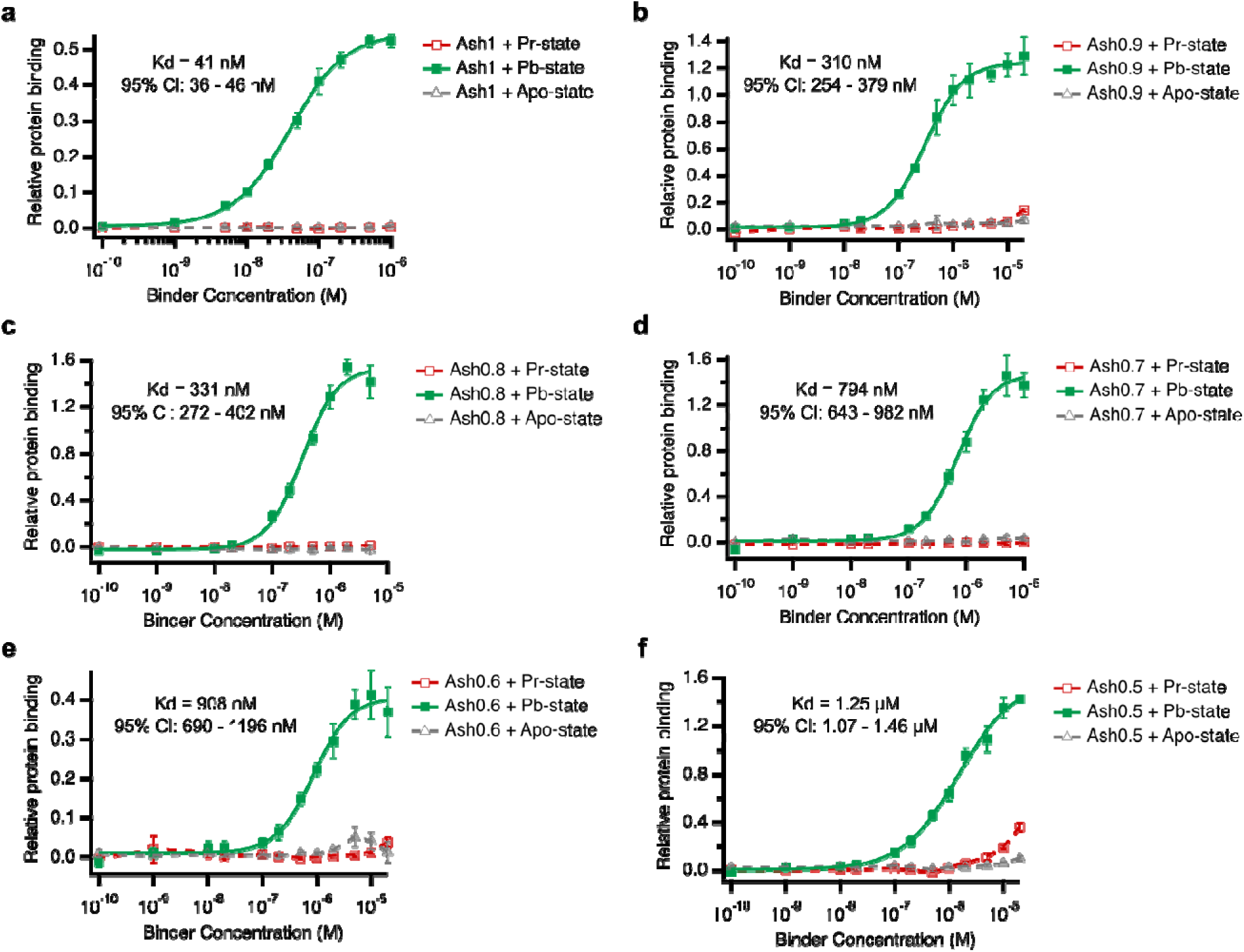
ELISA characterization of FenixS photo-state binders. ELISA results using purified, biotinylated holo- and apo-FenixS with purified binders Ash1 (**a**), Ash0.9 (**b**), Ash0.8 (**c**), Ash0.7 (**d**), Ash0.6 (**e**), and Ash0.5 (**f**). Dissociation constants (*K*_d_) and 95% confident intervals were estimated by fitting the data to a dose-response equation (f(x) = base + (max-base)/(1+*K*_d_/x)). n = 4 technical replicates, mean ± sd.

**Extended Data Fig. 6:**
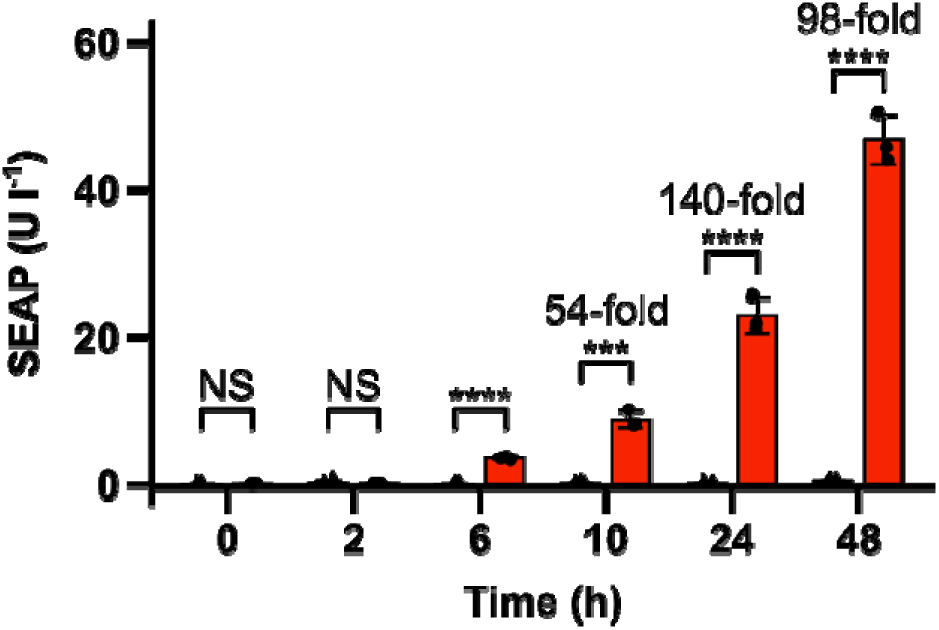
Dynamics of gene expression induced by FenixS-Ash1. SEAP expression at indicated time-points in CHO-K1 cells transiently transfected with the FenixS-FUS_N_-VP16-E-Ash1 construct and SEAP reporter plasmid. Cells were either kept in the dark or exposed to red light (680 nm, 0.59 mW cm^-2^).

**Extended Data Fig. 7:**
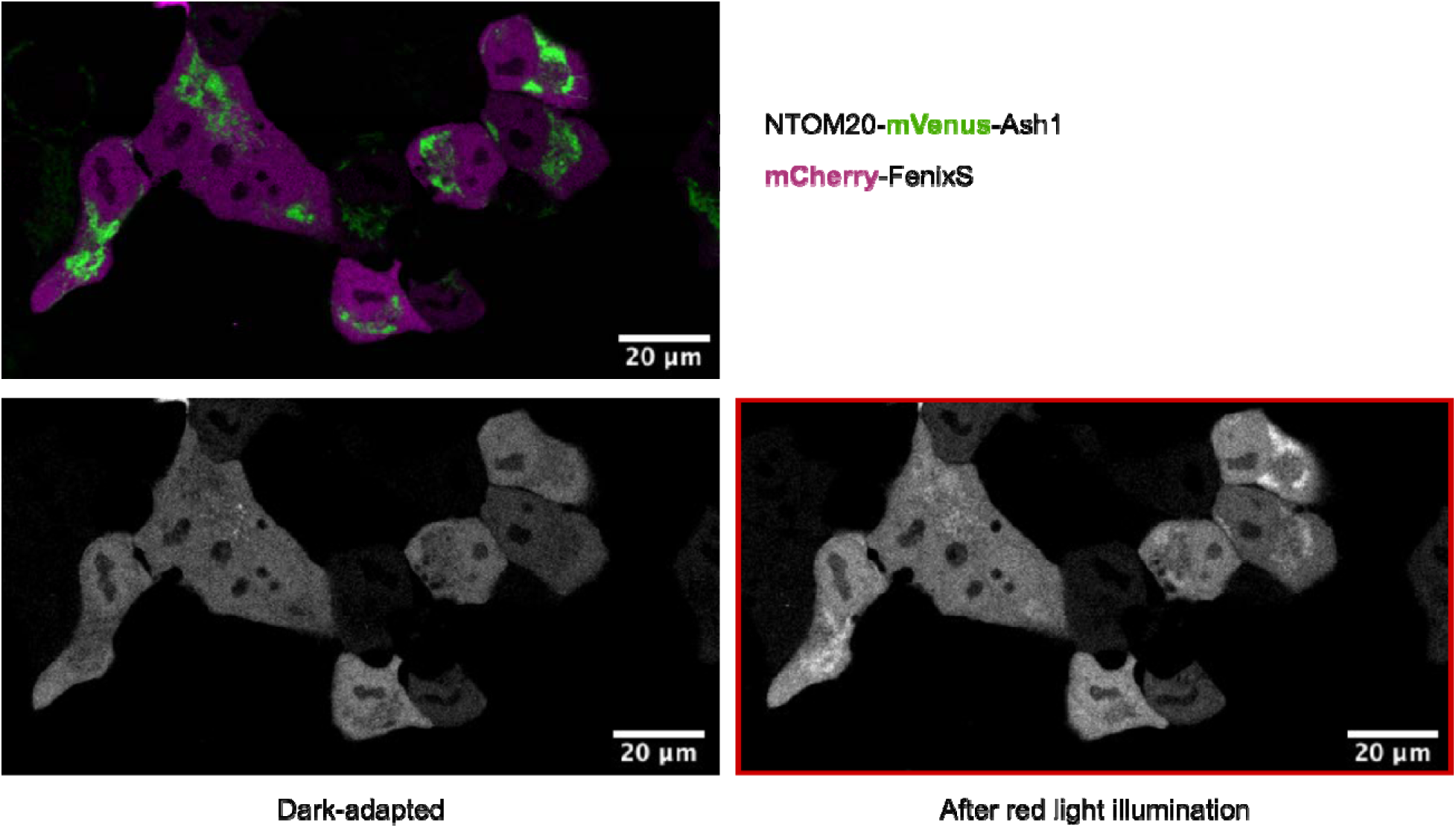
Light-controlled colocalization of FenixS with mitochondria-targeted Ash1 in Hela cells supplemented with 40 µM biliverdin. Representative confocal fluorescence images of mCherry-FenixS and NTOM20-mVenus-Ash1 in Hela cells at 40 µM supplemented biliverdin (BV). Top: Overlay of mitochondria-targeted mVenus-Ash1 and cytosolic mCherry-FenixS in the dark-adapted state. Bottom: mCherry fluorescence images in the dark-adapted state (left) and after red light illumination (680 nm) (right). Cells were imaged 24h post-transfection. Culture medium was supplemented with 40 µM BV 5 hours prior imaging. Scale bar, 20 µm.

## Notes

### Competing Interest Statement

The authors have declared no competing interest.

### Funding

- Natural Sciences and Engineering Research Council: RGPIN 175244

