## Supplementary material for "An ultra-low background far-red light-responsive optogenetic tool based on an engineered biliverdin-binding domain": Description of Addtional Movie Files

**Description of Additional Supplementary Files**

File name: Supplementary Movie 1

Description: Time-lapse imaging of FenixS-coated agarose beads incubated in 500 nM mCherry-Ash1 solution. Beads were imaged continuously using 20x air. Illumination was applied sequentially with red light (625 nm) followed by blue light (470 nm), as indicated by the text in the top-right corner. The timer shows minutes and seconds (mm:ss); scale bar = 40 μm.

File name: Supplementary Movie 2

Description: Time-lapse imaging of Hela cells expressing the mCherry-FenixS (shown) and mitochondria-localized NTOM20-mVenus-Ash1 (not shown). Cells were imaged using a 63x oil objective over iterative cycles of red (680 nm) and green (500 nm) illumination, as indicated by the text in the top-left corner. Each cycle consisted of 10 frames (270 s total) under 680 nm light followed by 15 frames (140 s total) under 500 nm light. Scale bar = 10 μm.

File name: Supplementary Movie 3

Description: Time-lapse imaging of HEK293T cells expressing the nuclear-localized NLS-FenixS-EGFP-FTH1 (not shown) and $FUS_N$-mCherry-Ash1 (shown). Cells were imaged continuously for 75 sec (30 frames) under red light (680 nm) using 63x oil objective. Scale bar: 10 μm.

File name: Supplementary Movie 4

Description: 3D reconstruction of a HEK293T cell expressing nuclear-localized NLS-FenixS-EGFP-FTH1 (green) and $FUS_N$-mCherry-Ash1 (magenta). Rendered using the 3D Viewer plugin. In the second half, the brightness threshold was increased to reveal condensates within the nucleus.
