## Supplementary Material for "An ultra-low background far-red light-responsive optogenetic tool based on an engineered biliverdin-binding domain"

### Supplementary Information

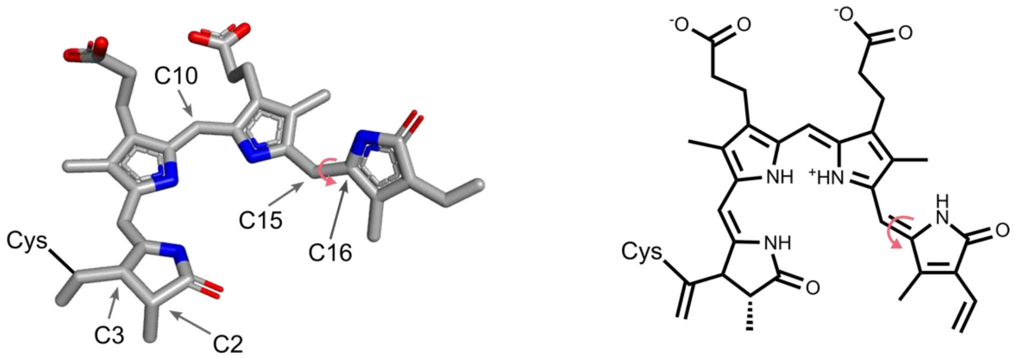

**Supplementary Figure 1: Biliverdin structure in miRFP670nano.**

Left: Stick representation of biliverdin bound to miRFP670nano (PDB ID: 6MGH[1]) rendered in Pymol. Right: Schematic skeleton structure. Carbon positions are labeled. Although FenixS was evolved from miRFP670nano, we hypothesize that its chromophore configuration more closely resembles that of canonical BV-binding CBCRs[2] (e.g. JSC1_58120g3), which typically feature a double bond between C2 and C3, a single bond between $C3^1$ and $C3^2$, and a cysteine attachment at either $C3^1$ or $C3^2$.

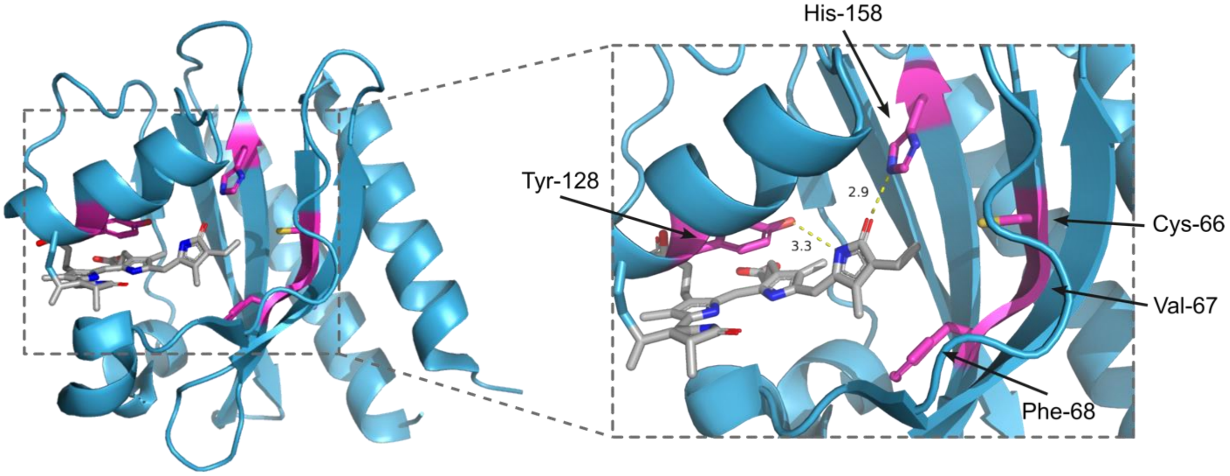

**Supplementary Figure 2: Structure of miRFP670nano.**

Crystal structure of miRFP670nano (PDB ID: 6MGH[1]). Amino acid residues surrounding the D-ring of the biliverdin chromophore that were introduced during the directed evolution of miRFP670nano are highlighted in pink.

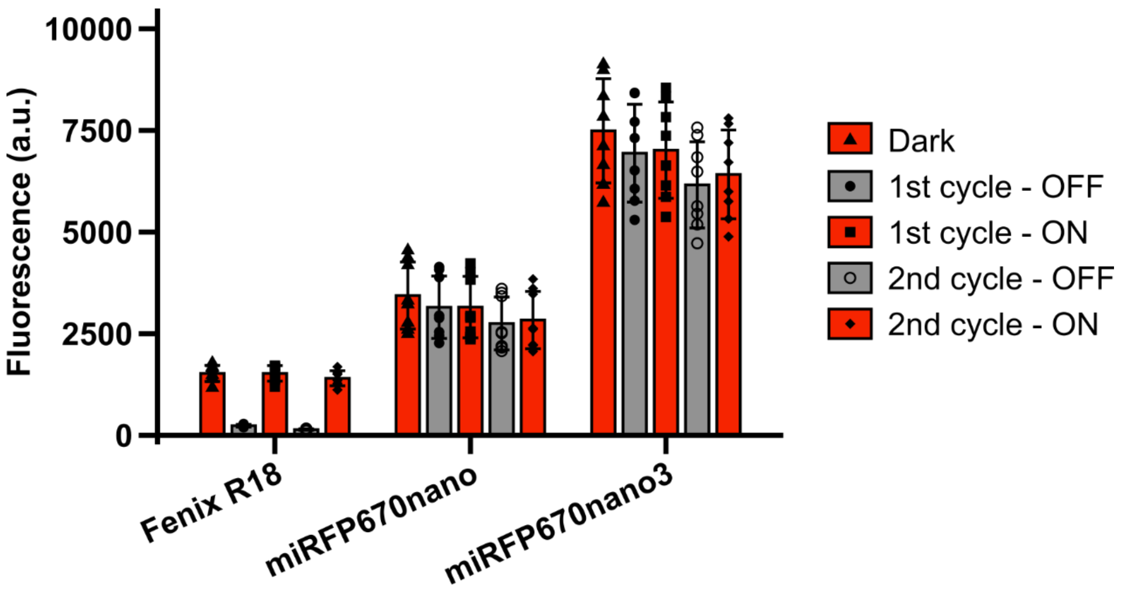

**Supplementary Figure 3: Comparison of photoswitchability between Fenix R18, miRFP670nano, and miRFP670nano3.**

Fluorescence and photoswitchability of Fenix R18, miRFP670nano, and miRFP670nano3 (crude protein extracts from *E. coli*). The OFF-state was produced with 660 nm illumination, the ON-state was produced with 530 nm illumination. Fluorescence of both states was detected at 670 nm with excitation at 647 nm. n = 8 biological replicates, mean ± sd.

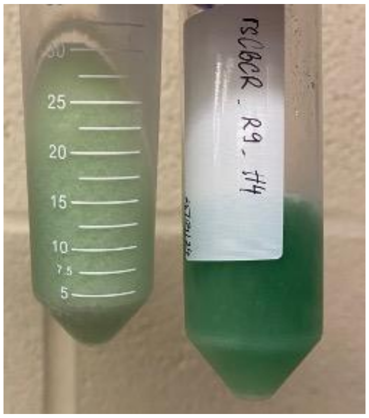

**Supplementary Figure 4: Effect of the R98S mutation on BV-binding efficiency in the Fenix R8 variant.**

Photographs of frozen *E. coli* cell pellets expressing Fenix R8 (left) and Fenix R8-R98S (right). The darker green color observed in the Fenix R8-R98S sample indicates a higher proportion of holo-protein, suggesting that the R98S mutation enhances BV-binding efficiency compared to the parental Fenix R8 variant.

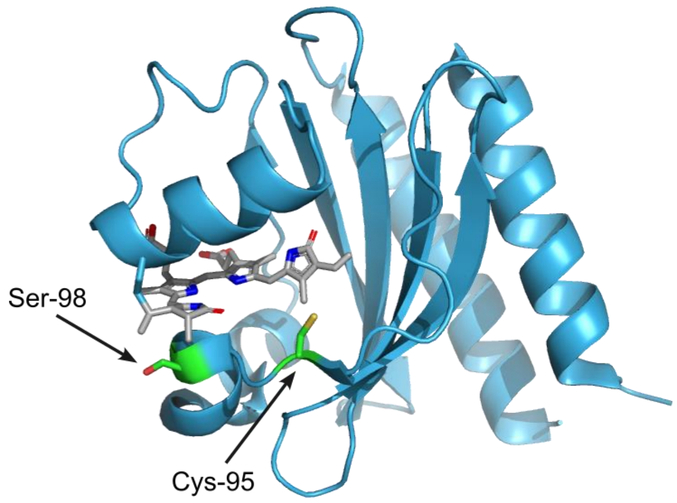

**Supplementary Figure 5: Positions of Ser98 and Cys95 mapped onto the miRFP670nano structure.**

Crystal structure of miRFP670nano (PDB ID: 6MGH[1]). Two key mutations introduced during the directed evolution of FenixS – R98S and V95C – are highlighted in green. Ser98 is located beneath the A-ring of the biliverdin chromophore, whereas Cys95 is located within a flexible loop positioned in front of the chromophore. This spatial arrangement of Cys95 suggests that formation of a C10-adduct photoproduct would likely induce substantial rearrangement of the chromophore-binding pocket.

| NpR3784 #s | 43 | 44 | 45 | 46 | 47 | 48 | 49 | 50 | 51 | 52 | 53 | 54 | 55 | 56 | 57 | 58 | 59 | 60 | 61 | 62 | 63 | 64 | 65 | 66 | 67 | 68 | 69 | 70 | 71 | 72 |
|---|---|---|---|---|---|---|---|---|---|---|---|---|---|---|---|---|---|---|---|---|---|---|---|---|---|---|---|---|---|---|
| FenixS | M | Y | Q | D | K | M | L | N | D | T | V | A | K | V | R | Q | I | L | Q | A | D | R | V | V | M | F | Q | F | E | E |
| miRFP670nano | M | N | L | D | K | M | L | N | T | T | V | T | E | V | R | Q | F | L | Q | V | D | R | V | C | V | F | Q | F | E | E |

| NpR3784 #s | 73 | 74 | 75 | 76 | 77 | 78 | 79 | 80 | 81 | 82 | 83 | 84 | 85 | 86 | 87 | 88 | 89 | 90 | 91 | 92 | 93 | 94 | 95 | 96 | 97 | 98 | 99 | 100 | 101 | 102 |
|---|---|---|---|---|---|---|---|---|---|---|---|---|---|---|---|---|---|---|---|---|---|---|---|---|---|---|---|---|---|---|
| FenixS | D | Y | S | G | E | V | V | V | E | A | V | D | G | R | W | S | S | I | L | G | T | Q | C | S | D | S | Y | F | M | E |
| miRFP670nano | D | Y | S | G | V | V | V | V | E | A | V | D | D | R | W | I | S | I | L | K | T | Q | V | R | D | R | Y | F | M | E |

| NpR3784 #s | 103 | 104 | 105 | 106 | 107 | 108 | 109 | 110 | 111 | 112 | 113 | 114 | 115 | 116 | 117 | 118 | 119 | 120 | 121 | 122 | 123 | 124 | 125 | 126 | 127 | 128 | 129 | 130 | 131 | 132 |
|---|---|---|---|---|---|---|---|---|---|---|---|---|---|---|---|---|---|---|---|---|---|---|---|---|---|---|---|---|---|---|
| FenixS | S | R | G | E | E | Y | A | R | G | R | Y | Q | A | I | A | D | I | Y | T | A | N | L | S | E | C | H | R | D | L | L |
| miRFP670nano | T | R | G | E | E | Y | S | H | G | R | Y | Q | A | I | A | D | I | Y | T | A | N | L | T | E | C | Y | R | D | L | L |

| NpR3784 #s | 133 | 134 | 135 | 136 | 137 | 138 | 139 | 140 | 141 | 142 | 143 | 144 | 145 | 146 | 147 | 148 | 149 | 150 | 151 | 152 | 153 | 154 | 155 | 156 | 157 | 158 | 159 | 160 | 161 | 162 |
|---|---|---|---|---|---|---|---|---|---|---|---|---|---|---|---|---|---|---|---|---|---|---|---|---|---|---|---|---|---|---|
| FenixS | A | Q | F | Q | V | R | A | V | L | A | V | P | I | L | Q | G | K | K | L | W | G | L | L | V | A | H | Q | L | A | A |
| miRFP670nano | T | Q | F | Q | V | R | A | I | L | A | V | P | I | L | Q | G | K | K | L | W | G | L | L | V | A | H | Q | L | A | A |

| NpR3784 #s | 163 | 164 | 165 | 166 | 167 | 168 | 169 | 170 | 171 | 172 | 173 | 174 | 175 | 176 | 177 | 178 | 179 | 180 | 181 | 182 | 183 | 184 | 185 | 186 | 187 | 188 |
|---|---|---|---|---|---|---|---|---|---|---|---|---|---|---|---|---|---|---|---|---|---|---|---|---|---|---|
| FenixS | P | R | Q | W | Q | S | W | E | I | D | F | L | K | Q | Q | A | V | A | V | G | I | A | I | Q | Q | I |
| miRFP670nano | P | R | Q | W | Q | T | W | E | I | D | F | L | K | Q | Q | A | V | V | V | G | I | A | I | Q | Q | S |

**Supplementary Figure 6: Sequence alignment of FenixS.**

Amino acid residues are indexed based on original template, NpR3784g1 (ref [3]). Mutations in FenixS relative to miRFP670nano are highlighted with an orange background. The R98S mutation is marked with a green background. The biliverdin-binding cysteine residue is indicated with a yellow background.

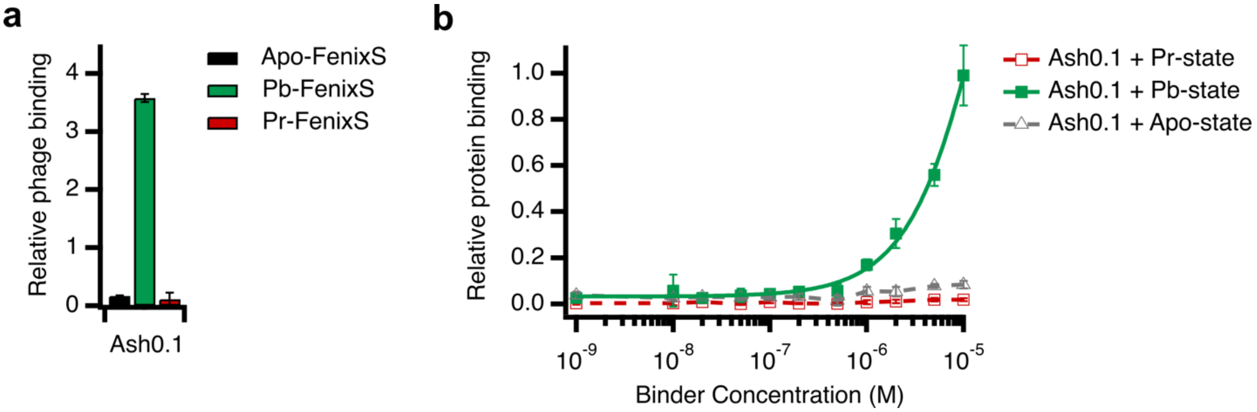

**Supplementary Figure 7: ELISA characterization of the FenixS photo-state binder, Ash0.1.**

**a,** pVIII-displayed phage ELISA results for Ash0.1 with biotinylated apo and holo-FenixS. n = 3 technical replicates, mean ± sd. Data represents results from two independent experiments. b, ELISA using purified Ash0.1 and biotinylated FenixS. n = 4 technical replicates, mean ± sd.

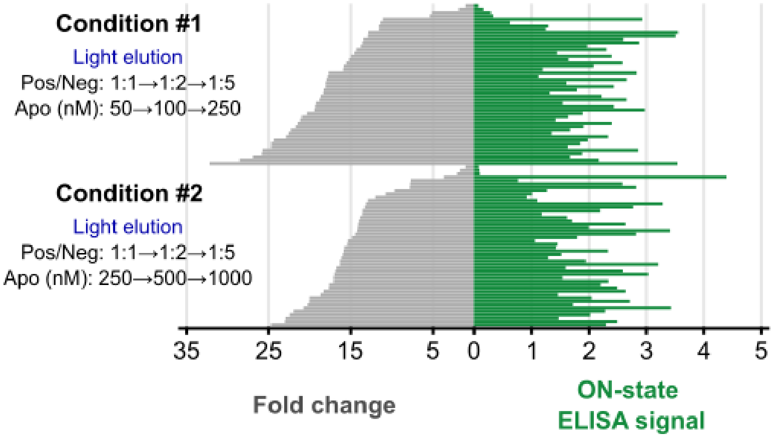

**Supplementary Figure 8: Single-clone phage ELISA during affinity maturation of Ash0.1.**

Single-clone ELISA results presented as butterfly plot from selection conditions #1 and #2 during the affinity maturation of Ash0.1. Each condition includes data from 48 individual clones. Detailed descriptions of the selection conditions are provided in the Methods section.

| Position | 1 | 2 | 3 | 4 | 5 | 6 | 7 | 8 | 9 | 10 | 11 | 12 | 13 | 14 | 15 | 16 | 17 | 18 | 19 | 20 | 21 | 22 | 23 | 24 | 25 | 26 | 27 | 28 |
|---|---|---|---|---|---|---|---|---|---|---|---|---|---|---|---|---|---|---|---|---|---|---|---|---|---|---|---|---|
| wt GA domain | T | I | D | Q | W | L | L | K | N | A | K | E | D | A | I | A | E | L | K | K | A | G | I | T | S | D | F | Y |
| Ash0.1 | T | I | D | Q | W | L | L | K | N | A | K | E | D | A | I | A | E | L | K | K | A | G | I | T | S | D | F | Y |
| Ash1.0 | T | I | D | Q | W | L | L | K | N | A | K | E | D | A | I | A | E | L | K | K | A | G | I | T | A | D | F | Y |
| Ash0.9 | T | I | D | Q | W | L | L | K | N | A | K | E | D | A | I | A | E | L | K | K | A | G | I | T | S | D | R | D |
| Ash0.8 | T | I | D | Q | W | L | L | K | N | A | K | E | D | A | I | A | E | L | K | K | A | G | I | T | S | D | R | E |
| Ash0.7 | T | I | D | Q | W | L | L | K | N | A | K | E | D | A | I | A | E | L | K | K | A | G | I | T | S | D | F | Y |
| Ash0.6 | T | I | D | Q | W | L | L | K | N | A | K | E | D | A | I | A | E | L | K | K | A | G | I | T | S | D | R | S |
| Ash0.5 | T | I | D | Q | W | L | L | K | N | A | K | E | D | A | I | A | E | L | K | K | A | G | I | T | C | D | F | Y |

| Position | 29 | 30 | 31 | 32 | 33 | 34 | 35 | 36 | 37 | 38 | 39 | 40 | 41 | 42 | 43 | 44 | 45 | 46 | 47 | 48 | 49 | 50 | 51 | 52 | 53 |
|---|---|---|---|---|---|---|---|---|---|---|---|---|---|---|---|---|---|---|---|---|---|---|---|---|---|
| wt GA domain | F | N | A | I | N | K | A | K | T | V | E | E | V | N | A | L | K | N | E | I | L | K | A | H | A |
| Ash0.1 | F | N | A | I | N | K | A | N | H | V | W | S | V | N | L | Y | K | N | E | I | L | K | A | H | A |
| Ash1.0 | F | N | V | I | N | K | A | N | Y | V | W | S | V | N | L | Y | K | N | D | I | L | K | A | H | A |
| Ash0.9 | F | N | V | I | N | K | A | N | H | V | W | S | V | N | L | Y | K | N | H | I | L | K | A | H | A |
| Ash0.8 | F | N | A | I | N | K | A | N | Y | V | W | S | V | N | V | Y | K | N | E | I | L | K | A | H | A |
| Ash0.7 | F | N | T | I | N | K | A | N | H | V | W | S | V | N | L | Y | K | N | D | I | L | K | A | H | A |
| Ash0.6 | F | N | T | I | N | K | A | H | H | V | W | S | V | N | L | Y | K | N | E | I | L | K | A | H | A |
| Ash0.5 | F | N | T | I | N | K | A | N | H | V | W | G | V | N | L | Y | K | N | D | I | L | K | A | H | A |

**Supplementary Figure 9: Amino acid sequences of binders targeting the Pg-state of FenixS.**

Amino acid sequences of binders obtained from naïve selection (Ash0.1) and subsequent affinity maturation (Ash0.5 – Ash1.0). Randomized residues are highlighted with background color corresponding to their amino acid identities.

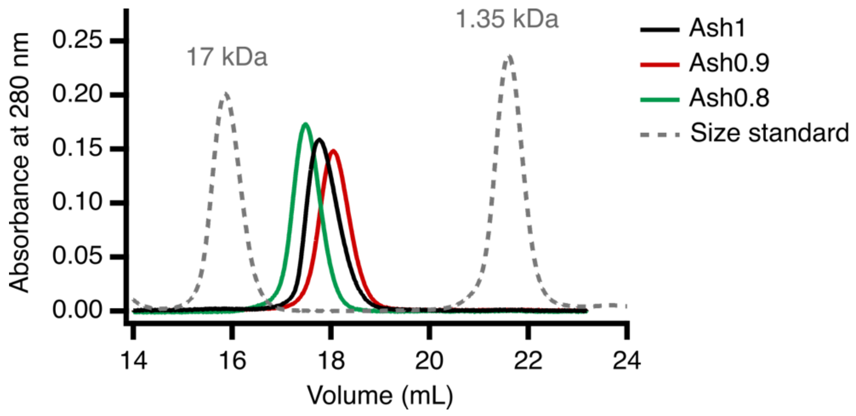

**Supplementary Figure 10: Size exclusion chromatography profiles of the purified binders.**

All proteins were injected at 50 μM. Molecular weight standard included myoglobin (17 kDa) and vitamin B12 (1.35 kDa) for reference.

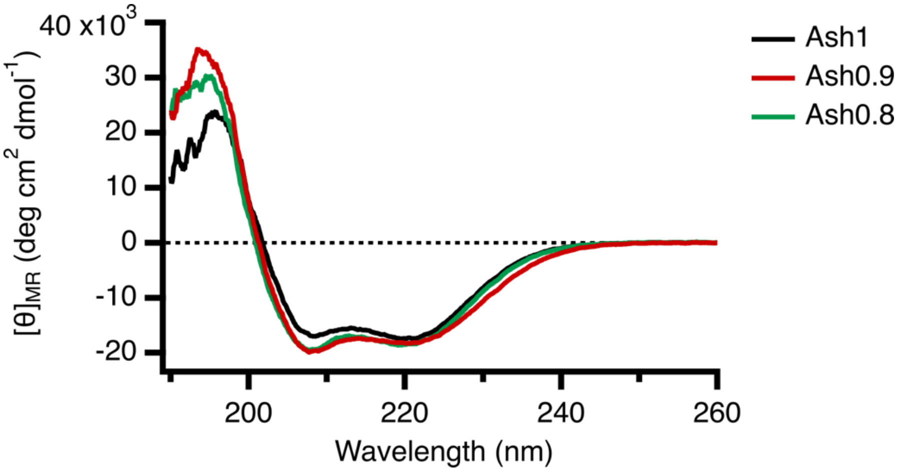

**Supplementary Figure 11: Circular dichroism (CD) spectra of the purified binders.**

All proteins were prepared at 50 μM in 10 mM sodium phosphate buffer and measured at room temperature.

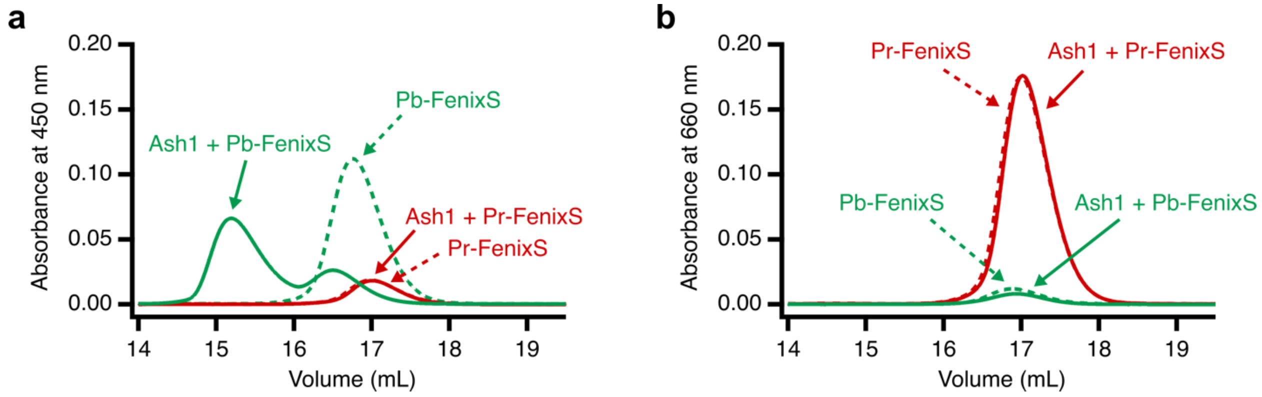

**Supplementary Figure 12: SEC profile of FenixS and Ash binding assay.**

Size-exclusion chromatography (SEC) elution profiles monitored by absorbance at 450 nm (**a**) and 660 nm (**b**). Curves represent dark-adapted and light-adapted FenixS, as well as mixtures of FenixS and Ash under the indicated conditions. Differences in elution profiles reflect light-dependent complex formation between FenixS and its photo-state-specific binder. In the Ash1 + Pb-FenixS trace, a small fraction of FenixS remains unbound to Ash1 (**a**). We hypothesize that this population did not switch to the Pb state, as indicated by residual absorbance at 660 nm. In contrast, the population that fully switched to the Pb state shows no absorbance at 660 nm (**b**), consistent with complete photo-conversion and binding.

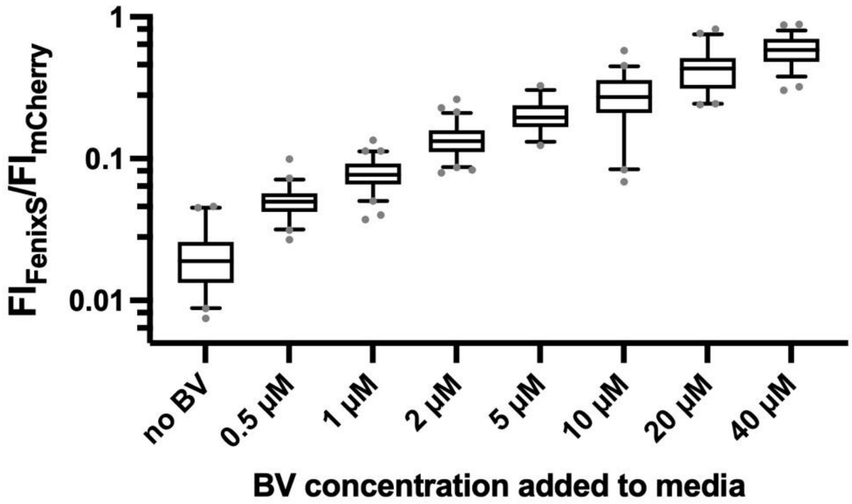

**Supplementary Figure 13: Quantification of holo-FenixS in Hela cells at varying concentration of supplemented BV.**

Fluorescence ratio of FenixS to mCherry in Hela cell at varying concentrations of supplemented biliverdin (BV). The fluorescence intensity of FenixS reflects the amount of holo-FenixS, while mCherry fluorescence correlates with the total expression level of both holo- and apo-FenixS. Thus, the FenixS-to-mCherry fluorescence ratio serves as proxy for the percentage of intracellular holo-FenixS. Cells were incubated with the indicated concentration of BV for 3 hours prior to imaging. Data represent n = 34 – 67 cells per condition.

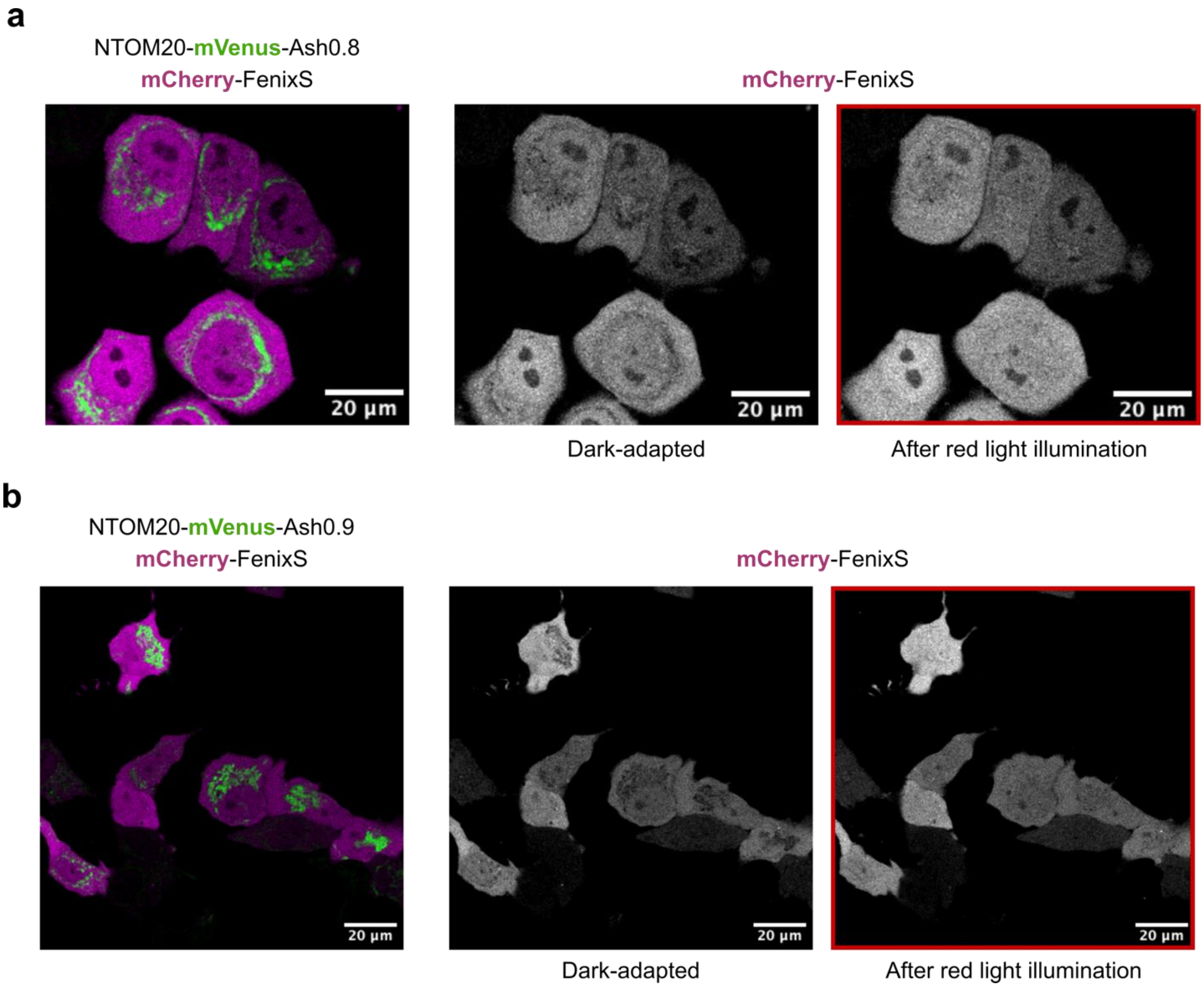

**Supplementary Figure 14: Light-controlled colocalization of FenixS with mitochondria-targeted Ash variants.**

Representative confocal fluorescence images of mCherry-FenixS and NTOM20-mVenus-Ash0.8 (**a**) or Ash0.9 (**b**) in Hela cells. Left panels show overlay images of mitochondria-targeted mVenus-binder (green) and cytosolic mCherry-FenixS (magenta) in the dark-adapted state. Right panels display mCherry fluorescence images in the dark-adapted state and after red light illumination (680 nm). Scale bar, 20 µm.

#### Supplementary Table 1. Plasmids designed and used in this study

| Plasmid name | Plasmid vector | Expression in | Promoter | Features | Used in | Reference |
|---|---|---|---|---|---|---|
| P1 | pBAD/HisB | Bacteria | araBAD | miRFP670nano-His6::HO1 | Extended Fig. 1; Fig. S3 | This work |
| P2 | pBAD/HisB | Bacteria | araBAD | miRFP670nano3-His6::HO1 | Fig. S3 | This work |
| P3 | pBAD/HisB | Bacteria | araBAD | Fenix (R0-R18)-His6::HO1 | Fig. 1b; Extended Fig. 1,2; Fig. S3, S4 | This work |
| P4 | pBAD/HisB | Bacteria | araBAD | FenixS-His6::HO1 | Fig. 1e-f, 2e; Extended Fig. 3, 4; Fig. S12 | This work |
| P5 | pET24b | Bacteria | T7 | FenixS-His6 | Fig. 1d | This work |
| P6 | pET24b | Bacteria | T7 | miRFP670nano-His6 | Fig. 1d | This work |
| P7 | pET24b | Bacteria | T7 | AM1_C0023g2 S334G-His6 | Fig. 1d | Jaewan. J. *et. al.* 2023[4] |
| P8 | pBAD/HisB | Bacteria | araBAD | Avi-FenixS-His6::HO1 | Fig. 2c-d, 3b; Extended Fig. 5; Fig. S8 | This work |
| P9 | pET24b | Bacteria | T7 | Avi-FenixS-His6 | Fig. 2c; Extended Fig. 5 | This work |
| P10 | p8 | Bacteria | TAC | stII (secretion signal)-FLAG-GA library-truncated P8 | Fig. S7a | Reis, J. *et al.* 2018[5] |
| P11 | p3 | Bacteria | TAC | stII (secretion signal)-FLAG-GA library-DKTHTCGRP (dimerization sequence)-Cterm. P3 | Fig. 2c; Fig. S8 | This work |

| | | | | | | |
|---|---|---|---|---|---|---|
| P12 | pET24b | Bacteria | T7 | FLAG-Ash0.1-His6 | Fig. S7b | This work |
| P13 | pET24b | Bacteria | T7 | FLAG-Ash1-His6 | Fig. 2d-e; Extended Fig. 5a; Fig. S10, S11, S12, S15 | This work |
| P14 | pET24b | Bacteria | T7 | FLAG-Ash0.8-His6 | Fig. 2d; Extended Fig. 5c; Fig. S10, S11 | This work |
| P15 | pET24b | Bacteria | T7 | FLAG-Ash0.9-His6 | Fig. 2d; Extended Fig. 5b; Fig. S10, S11 | This work |
| P16 | pET24b | Bacteria | T7 | FLAG-Ash0.7-His6 | Extended Fig. 5d | This work |
| P17 | pET24b | Bacteria | T7 | FLAG-Ash0.6-His6 | Extended Fig. 5e | This work |
| P18 | pET24b | Bacteria | T7 | FLAG-Ash0.5-His6 | Extended Fig. 5f | This work |
| P19 | pET24b | Bacteria | T7 | mCherry-Ash1-His6 | Fig. 3b | This work |
| P20 | pTRIEX | Mammalian | CMV | mCherry-FenixS | Fig. 5b-e; Extended Fig. 7; Fig. S13. S14 | This work |
| P21 | pTRIEX | Mammalian | CMV | NTOM20-mVenus-Ash1 | Fig. 5b-e; Extended Fig. 7 | This work |
| P22 | pTRIEX | Mammalian | CMV | NTOM20-mVenus-Ash0.8 | Fig. S14 | This work |
| P23 | pTRIEX | Mammalian | CMV | NTOM20-mVenus-Ash0.9 | Fig. S14 | This work |
| P24 | pHR | Mammalian | SFFV | FUSN-mCherry-Ash1 | Fig. 6b-f | This work |
| P25 | pHR | Mammalian | SFFV | NLS-FenixS-EGFP-FTH1 | Fig. 6b-f | This work |

| | | | | | | |
|---|---|---|---|---|---|---|
| P26 | pWW035 | Mammalian | SV40 | E-VP16 | Fig. 4b | Muller, K. et al. 2013[6] |
| P27 | pKT214 | Mammalian | SV40 | Amg2-VP16-NLS-IRES-E-BAmRed1.4 | N.A. | Jang, J. et al. 2023[4] |
| P28 | pKT753 | Mammalian | SV40 | Amg2-FUS-VP16-NLS-IRES-E-BAmRed1.4 | N.A. | Jang, J. et al. 2023[4] |
| P29 | pKM081 | Mammalian | CMVmin | SEAP | Fig. 4b-d; Extended Fig. 6 | Muller, K. et al. 2013[6] |
| P30 | pKT214 | Mammalian | SV40 | FenixS-VP16-NLS-IRES-E-Ash1 | Fig. 4b | This work |
| P31 | pKT753 | Mammalian | SV40 | FenixS-FUS-VP16-NLS-IRES-E-Ash1 | Fig. 4b-d; Extended Fig. 6 | This work |

**Supplementary Note: DNA sequence of plasmid designed and used in this study**

**P1 miRFP670nano-His6**

ATGAACCTGGACAAGATGCTGAATACCACAGTAACAGAGGTGCGGCAGTTCCTGCAGGTGG
ACAGAGTGTGCGTGTTCCAGTTTGAGGAGGATTATAGCGGAGTGGTGGTGGAGGCCG
TGGACGATAGGTGGATCTCCATCCTGAAGACCCAGGTGCGGGATAGATACTTCATGGAGAC
AAGGGGCGAGGAGTATTCTCACGGCCGCTACCAGGCCATCGCCGACATCTACACCGCAAA
CCTGACAGAGTGCTACAGGGATCTGCTGACACAGTTTCAGGTGAGAGCAATCCTGGCCGT
GCCCATCCTGCAGGGCAAGAAGCTGTGGGGCCTGTTGGTGGCACACCAGCTGGCGGCCC
CTAGACAGTGGCAGACCTGGGAGATCGACTTTCTGAAGCAGCAGGCCGTGGTGGTGGGCA
TCGCCATCCAGCAGAGCGGGAGCTCTGGAAAGCTTCATCATCACCATCACCATTGA

**P2 miRFP670nano3-His6**

ATGAACCTGGACAAGATGCTGAACACCACCGTGACCGAGGTGCGCAAGTTCCTGCAAGCG
GACAGAGTGTGCGTGTTCAAGTTCGAGGAAGATTACTCCGGCACCGTCTCGCACGAAGCC
GTGGACGACAGATGGATTAGCATCCTGAAGACCCAGGTGCAGGACAGATACTTCATGGAAA
CCAGAGGCGAGGAATACGTCCACGGCAGATACCAGGCCATCGCCGACATCTACACAGCCA
ATCTGGTCGAGTGCTACAGAGACCTGCTGATCGAGTTTCAGGTGCGGGCCATTCTGGCTGT
CCCCATCCTGCAAGGCAAGAAGCTGTGGGGCCTGCTGGTGGCCCACCAACTGGCCGGCC
CTCGGGAGTGGCAGACCTGGGAAATCGACTTCCTGAAACAGCAAGCCGTGGTGATGGGCA
TCGCCATCCAGCAGAGCGGGAGCTCTGGAAAGCTTCATCATCACCATCACCATTGA

**P3 Fenix R18-His6**

ATGTACCAGGACAAGATGTTGAATGACACAGTAGCAAAGGTGCGACAGATCCTGCAGGCGG
ACAGGGTGGTTATGTTCCAATTCGAGGAGGATTACAGCGGAGAGGTGGTGGTGGAGGCCG
TGGACGGTAGGTGGAGCTCCATCCTGGGGACCCAGTGTAGTGATAGATACTTCATGGAGTC
AAGGGGCGAGGAGTATGCTCGCGGCCGCTACCAGGCCATCGCCGACATCTACACCGCTAA
CCTGTCAGAGTGCCACAGGGATCTGCTGGCGCAGTTTCAGGTGAGAGCGGTCCTGGCCGT
GCCCATCCTGCAGGGCAAGAAGCTGTGGGGCCTGTTGGTGGCACACCAGCTGGCGGCCC
CTAGACAGTGGCAGTCCTGGGAGATCGACTTTCTGAAGCAGCAGGCCGTGGCGGTGGGC
ATAGCCATCCAACAGATCGGGAGCTCTGGAAAGCTTCATCATCACCATCACCATTGA

**P4 (pBAD) FenixS-His6**

ATGTACCAGGACAAGATGTTGAATGACACAGTAGCAAAGGTGCGACAGATCCTGCAGGCGG
ACAGGGTGGTTATGTTCCAATTCGAGGAGGATTACAGCGGAGAGGTGGTGGTGGAGGCCG
TGGACGGTAGGTGGAGCTCCATCCTGGGGACCCAGTGTAGTGATAGTTACTTCATGGAGTC

AAGGGGCGAGGAGTATGCTCGCGGCCGCTACCAGGCCATCGCCGACATCTACACCGCTAA
CCTGTCAGAGTGCCACAGGGATCTGCTGGCGCAGTTTCAGGTGAGAGCGGTCCTGGCCGT
GCCCATCCTGCAGGGCAAGAAGCTGTGGGGCCTGTTGGTGGCACACCAGCTGGCGGCCC
CTAGACAGTGGCAGTCCTGGGAGATCGACTTTCTGAAGCAGCAGGCCGTGGCGGTGGGC
ATAGCCATCCAACAGATCGGGAGCTCTGGAAAGCTTCATCATCACCATCACCATTGA

**P5 (pET24b) FenixS-His6**

ATGTACCAGGACAAGATGTTGAATGACACAGTAGCAAAGGTGCGACAGATCCTGCAGGCGG
ACAGGGTGGTTATGTTCCAATTCGAGGAGGATTACAGCGGAGAGGTGGTGGTGGAGGCCG
TGGACGGTAGGTGGAGCTCCATCCTGGGGACCCAGTGTAGTGATAGTTACTTCATGGAGTC
AAGGGGCGAGGAGTATGCTCGCGGCCGCTACCAGGCCATCGCCGACATCTACACCGCTAA
CCTGTCAGAGTGCCACAGGGATCTGCTGGCGCAGTTTCAGGTGAGAGCGGTCCTGGCCGT
GCCCATCCTGCAGGGCAAGAAGCTGTGGGGCCTGTTGGTGGCACACCAGCTGGCGGCCC
CTAGACAGTGGCAGTCCTGGGAGATCGACTTTCTGAAGCAGCAGGCCGTGGCGGTGGGC
ATAGCCATCCAACAGATCGGGAGCTCTGGAAAGCTTCACCACCACCACCACCACTGA

**P6 (pET24b) miRFP670nano-His6**

ATGAAACTGGCCAACCTGGACAAGATGCTGAATACCACAGTAACAGAGGTGCGGCAGTTCC
TGCAGGTGGACAGAGTGTGCGTGTTCCAGTTTGAGGAGGATTATAGCGGAGTGGTGGTGG
TGGAGGCCGTGGACGATAGGTGGATCTCCATCCTGAAGACCCAGGTGCGGGATAGATACTT
CATGGAGACAAGGGGCGAGGAGTATTCTCACGGCCGCTACCAGGCCATCGCCGACATCTA
CACCGCAAACCTGACAGAGTGCTACAGGGATCTGCTGACACAGTTTCAGGTGAGAGCAATC
CTGGCCGTGCCCATCCTGCAGGGCAAGAAGCTGTGGGGCCTGTTGGTGGCACACCAGCT
GGCGGCCCCTAGACAGTGGCAGACCTGGGAGATCGACTTTCTGAAGCAGCAGGCCGTGG
TGGTGGGCATCGCCATCCAGCAGAGCGGGAGCTCTGGACTCGAGCACCACCACCACCAC
CACTGA

**P7 (pET24b) AM1_C0023g2 S334G-His6**

ATGAAACTGGCCAAAAGCCTGGATATCGAAGATATCTTCGGTGCGACCACCCAGGATGTTC
GTGAATCTCTGGAATGTGATCGTGTTGTGATCTACCAGTTTTGGCCGGATTGGTCTGGTGAA
TTCCTGGTTGAATCCACCGCGCCGGGTCTGATCCCGCTGTCTGAACTGGATGTTCCGATGA
CCTGGCAGGATACCTACCTGCAGGAAAACCAGGGCGGTAAATTCAAAGATAACGCACCGAC
CGTTGTTGCTGATATTTATCAGCAGGGTTACACTGATTGTCACCTGGAAATCCTGGAATGGTT
TGATATTCGTGCTTACATGGTTGTGCCGGTTTTCATTGGTAAAACCTTATGGGGTCTGTTAGC
TGCTTATCAGCTGAACCATCCGCGTCAGTGGCAGAAAGTTGAACTGTATCTGCTGAAACAG

GCGGGTGCTCAGCTGGGTGTTGCACTGCAGCAGGCTGAACTGCTGAACCAGAAGCTCGA
GCACCACCACCACCACCACTGA

**P8 (pBAD) Avi tag-FenixS-His6**

ATGAAAGGTCTGAACGACATCTTCGAGGCTCAGAAAATCGAATGGCACGAAGGATCCAGCG
GAATGTACCAGGACAAGATGTTGAATGACACAGTAGCAAAGGTGCGACAGATCCTGCAGGC
GGACAGGGTGGTTATGTTCCAATTCGAGGAGGATTACAGCGGAGAGGTGGTGGAGGC
CGTGGACGGTAGGTGGAGCTCCATCCTGGGGACCCAGTGTAGTGATAGTTACTTCATGGAG
TCAAGGGGCGAGGAGTATGCTCGCGGCCGCTACCAGGCCATCGCCGACATCTACACCGCT
AACCTGTCAGAGTGCCACAGGGATCTGCTGGCGCAGTTTCAGGTGAGAGCGGTCCTGGCC
GTGCCCATCCTGCAGGGCAAGAAGCTGTGGGGCCTGTTGGTGGCACACCAGCTGGCGGC
CCCTAGACAGTGGCAGTCCTGGGAGATCGACTTTCTGAAGCAGCAGGCCGTGGCGGTGG
GCATAGCCATCCAACAGATCGGGAGCTCTGGAAAGCTTCATCATCACCATCACCATTGA

**P9 (pET24b) Avi tag-FenixS-His6**

ATGAAAGGTCTGAACGACATCTTCGAGGCTCAGAAAATCGAATGGCACGAAGGATCCAGCG
GAATGTACCAGGACAAGATGTTGAATGACACAGTAGCAAAGGTGCGACAGATCCTGCAGGC
GGACAGGGTGGTTATGTTCCAATTCGAGGAGGATTACAGCGGAGAGGTGGTGGAGGC
CGTGGACGGTAGGTGGAGCTCCATCCTGGGGACCCAGTGTAGTGATAGTTACTTCATGGAG
TCAAGGGGCGAGGAGTATGCTCGCGGCCGCTACCAGGCCATCGCCGACATCTACACCGCT
AACCTGTCAGAGTGCCACAGGGATCTGCTGGCGCAGTTTCAGGTGAGAGCGGTCCTGGCC
GTGCCCATCCTGCAGGGCAAGAAGCTGTGGGGCCTGTTGGTGGCACACCAGCTGGCGGC
CCCTAGACAGTGGCAGTCCTGGGAGATCGACTTTCTGAAGCAGCAGGCCGTGGCGGTGG
GCATAGCCATCCAACAGATCGGGAGCTCTGGAAAGCTTCACCACCACCACCACCACTGA

**P11 stII (secretion signal)-FLAG-GA library (Ash0.1)-Dimerization sequence-Cterm. P3**

ATGAAAAAGAATATCGCATTTCTTCTTGCATCTATGTTCGTTTTTTCTATTGCTACAAATGCCTA
TGCATCCGATTATAAAGATGATGATGATAAAGGCGGATCCACGATTGACCAGTGGCTGCTGA
AAAACGCGAAAGAAGATGCTATTGCAGAACTGAAAAAGGCTGGTATCACCTCTGACTTTTAC
TTCAACGCGATCAATAAAGCGAATCATGTGTGGTCTGTTAACTTGTATAAGAACGAGATCCTG
AAAGCTCACGCCGGGAGCTCTGGAGACAAAACTCACACATGCGGCCGGCCCTCTGGTTCC
GGTGATTTTGATTATGAAAAGATGGCAAACGCTAATAAGGGGGCTATGACCGAAAATGCCGA
TGAAAACGCGCTACAGTCTGACGCTAAAGGCAAACTTGATTCTGTCGCTACTGATTACGGTG
CTGCTATCGATGGTTTCATTGGTGACGTTTCCGGCCTTGCTAATGGTAATGGTGCTACTGGT
GATTTTGCTGGCTCTAATTCCCAAATGGCTCAAGTCGGTGACGGTGATAATTCACCTTTAATG

AATAATTTCCGTCAATATTTACCTTCCCTCCCTCAATCGGTTGAATGTCGCCCTTTTGTCTTTA
GCGCTGGTAAACCATATGAATTTTCTATTGATTGTGACAAAATAAACTTATTCCGTGGTGTCTT
TGCGTTTCTTTTATATGTTGCCACCTTTATGTATGTATTTTCTACGTTTGCTAACATACTGCGTA
ATAAGGAGTCTTAA

**P12 FLAG-Ash0.1-His6**

ATGAAAGATTATAAAGATGATGATGATAAAGGCGGATCCACGATTGACCAGTGGCTGCTGAA
AAACGCGAAAGAAGATGCTATTGCAGAACTGAAAAAGGCTGGTATCACCTCTGACTTTTACT
TCAACGCGATCAATAAAGCGAATCATGTGTGGTCTGTTAACTTGTATAAGAACGAGATCCTGA
AAGCTCACGCCGGGAGCTCTGGAAAGCTTCACCACCACCACCACCACTGA

**P13 FLAG-Ash1-His6**

ATGAAAGATTATAAAGATGATGATGATAAAGGCGGATCCACGATTGACCAGTGGCTGCTGAA
AAACGCGAAAGAAGATGCTATTGCAGAACTGAAAAAGGCTGGTATCACCGCTGACTTTTACT
TCAACGTGATCAATAAAGCGAACTATGTGTGGTCTGTTAACTTGTATAAGAACGATATCCTGA
AAGCTCACGCCGGGAGCTCTGGAAAGCTTCACCACCACCACCACCACTGA

**P14 FLAG-Ash0.8-His6**

ATGAAAGATTATAAAGATGATGATGATAAAGGCGGATCCACGATTGACCAGTGGCTGCTGAA
AAACGCGAAAGAAGATGCTATTGCAGAACTGAAAAAGGCTGGTATCACCTCTGACCGTGAA
TTCAACGCGATCAATAAAGCGAATTATGTGTGGTCTGTTAACGTGTACAAGAACGAGATCCT
GAAAGCTCACGCCGGGAGCTCTGGAAAGCTTCACCACCACCACCACCACTGA

**P15 FLAG-Ash0.9-His6**

ATGAAAGATTATAAAGATGATGATGATAAAGGCGGATCCACGATTGACCAGTGGCTGCTGAA
AAACGCGAAAGAAGATGCTATTGCAGAACTGAAAAAGGCTGGTATCACCTCAGACCGGGAC
TTCAACGTGATCAATAAAGCGAATCATGTGTGGTCTGTTAACTTGTATAAGAACCACATCCTG
AAAGCTCACGCCGGGAGCTCTGGAAAGCTTCACCACCACCACCACCACTGA

**P16 FLAG-Ash0.7-His6**

ATGAAAGATTATAAAGATGATGATGATAAAGGCGGATCCACGATTGACCAGTGGCTGCTGAA
AAACGCGAAAGAAGATGCTATTGCAGAACTGAAAAAGGCTGGTATCACCTCGGACTTTTACT
TCAACACGATCAATAAAGCGAATCATGTGTGGTCTGTTAACTTGTATAAGAACGATATCCTGA
AAGCTCACGCCGGGAGCTCTGGAAAGCTTCACCACCACCACCACCACTGA

**P17 FLAG-Ash0.6-His6**

ATGAAAGATTATAAAGATGATGATGATAAAGGCGGATCCACGATTGACCAGTGGCTGCTGAA
AAACGCGAAAGAAGATGCTATTGCAGAACTGAAAAAGGCTGGTATCACCTCTGACCGTTCTT
TCAACACGATCAATAAAGCGCATCATGTGTGGTCTGTTAACTTGTATAAGAACGAGATCCTGA
AAGCTCACGCCGGGAGCTCTGGAAAGCTTCACCACCACCACCACCACTGA

**P18 FLAG-Ash0.5-His6**

ATGAAAGATTATAAAGATGATGATGATAAAGGCGGATCCACGATTGACCAGTGGCTGCTGAA
AAACGCGAAAGAAGATGCTATTGCAGAACTGAAAAAGGCTGGTATCACCTGCGACTTTTATT
TCAACACGATCAATAAAGCGAATCACGTGTGGGGTGTTAACTTGTATAAGAACGATATCCTGA
AAGCTCACGCCGGGAGCTCTGGAAAGCTTCACCACCACCACCACCACTGA

**P19 mCherry-Ash1-His6**

ATGGTGAGCAAGGGCGAGGAGGATAACATGGCCATCATCAAGGAGTTCATGCGCTTCAAGG
TGCACATGGAGGGCTCCGTGAACGGCCACGAGTTCGAGATCGAGGGCGAGGGCGAGGGC
CGCCCCTACGAGGGCACCCAGACCGCCAAGCTGAAGGTGACCAAGGGTGGCCCCCTGCC
CTTCGCCTGGGACATCCTGTCCCCTCAGTTCATGTACGGCTCCAAGGCCTACGTGAAGCAC
CCCGCCGACATCCCCGACTACTTGAAGCTGTCCTTCCCCGAGGGCTTCAAGTGGGAGCGC
GTGATGAACTTCGAGGACGGCGGCGTGGTGACCGTGACCCAGGACTCCTCCCTGCAGGA
CGGCGAGTTCATCTACAAGGTGAAGCTGCGCGGCACCAACTTCCCCTCCGACGGCCCTGT
AATGCAGAAGAAGACTATGGGCTGGGAGGCCTCCTCCGAGCGGATGTACCCCGAGGACGG
CGCCCTGAAGGGCGAGATCAAGCAGAGGCTGAAGCTGAAGGACGGCGGCCACTACGACG
CTGAGGTCAAGACCACCTACAAGGCCAAGAAGCCCGTGCAGCTGCCCGGCGCCTACAACG
TCAACATCAAGTTGGACATCACCTCCCACAACGAGGACTACACCATCGTGGAACAGTACGA
ACGCGCCGAGGGCCGCCACTCCACCGGCGGCATGGACGAGCTGTACAAGGGCGGATCCA
CGATTGACCAGTGGCTGCTGAAAAACGCGAAAGAAGATGCTATTGCAGAACTGAAAAAGGC
TGGTATCACCGCTGACTTTTACTTCAACGTGATCAATAAAGCGAACTATGTGTGGTCTGTTAA
CTTGTATAAGAACGATATCCTGAAAGCTCACGCCGGGAGCTCTGGAAAGCTTCACCACCAC
CACCACCACTGA

**P20 mCherry-FenixS**

ATGGTGAGCAAGGGCGAGGAGGATAACATGGCCATCATCAAGGAGTTCATGCGCTTCAAGG
TGCACATGGAGGGCTCCGTGAACGGCCACGAGTTCGAGATCGAGGGCGAGGGCGAGGGC
CGCCCCTACGAGGGCACCCAGACCGCCAAGCTGAAGGTGACCAAGGGTGGCCCCCTGCC
CTTCGCCTGGGACATCCTGTCCCCTCAGTTCATGTACGGCTCCAAGGCCTACGTGAAGCAC
CCCGCCGACATCCCCGACTACTTGAAGCTGTCCTTCCCCGAGGGCTTCAAGTGGGAGCGC

GTGATGAACTTCGAGGACGGCGGCGTGGTGACCGTGACCCAGGACTCCTCCCTGCAGGA
CGGCGAGTTCATCTACAAGGTGAAGCTGCGCGGCACCAACTTCCCCTCCGACGGCCCTGT
AATGCAGAAGAAGACTATGGGCTGGGAGGCCTCCTCCGAGCGGATGTACCCCGAGGACGG
CGCCCTGAAGGGCGAGATCAAGCAGAGGCTGAAGCTGAAGGACGGCGGCCACTACGACG
CTGAGGTCAAGACCACCTACAAGGCCAAGAAGCCCGTGCAGCTGCCCGGCGCCTACAACG
TCAACATCAAGTTGGACATCACCTCCCACAACGAGGACTACACCATCGTGGAACAGTACGA
ACGCGCCGAGGGCCGCCACTCCACCGGCGGCATGGACGAGCTGTACAAGGGTTCTGGAT
CCGGTTCTGGCTCAATGTACCAGGACAAGATGTTGAATGACACAGTAGCAAAGGTGCGACA
GATCCTGCAGGCGGACAGGGTGGTTATGTTCCAATTCGAGGAGGATTACAGCGGAGAGGT
GGTGGTGGAGGCCGTGGACGGTAGGTGGAGCTCCATCCTGGGGACCCAGTGTAGTGATA
GTTACTTCATGGAGTCAAGGGGCGAGGAGTATGCTCGCGGCCGCTACCAGGCCATCGCCG
ACATCTACACCGCTAACCTGTCAGAGTGCCACAGGGATCTGCTGGCGCAGTTTCAGGTGAG
AGCGGTCCTGGCCGTGCCCATCCTGCAGGGCAAGAAGCTGTGGGGCCTGTTGGTGGCAC
ACCAGCTGGCGGCCCCTAGACAGTGGCAGTCCTGGGAGATCGACTTTCTGAAGCAGCAGG
CCGTGGCGGTGGGCATAGCCATCCAACAGATCTAA

**P21 NTOM20-mVenus-Ash1-His6**

ATGGTGGGCCGCAACAGCGCGATTGCAGCTGGAGTGTGCGGCGCGCTGTTCATTGGCTAT
TGCATTTACTTTGATCGCAAACGCCGCAGCGATCCGAACTTTAAATCTAGAGTGAGCAAGGG
CGAGGAGCTGTTCACCGGGGTGGTGCCCATCCTGGTCGAGCTGGACGGCGACGTAAACG
GCCACAAGTTCAGCGTGTCCGGCGAGGGCGAGGGCGATGCCACCTACGGCAAGCTGACC
CTGAAGTTCATCAGCACCACCGGCAAGCTGCCCGTGCCCTGGCCCACCCTCGTGACCACC
CTCGGCTACGGCCTGCAGTGCTTCGCCCGCTACCCCGACCACATGAAGCAGCACGACTTC
TTCAAGTCCGCCATGCCCGAAGGCTACGTCCAGGAGCGCACCATCTTCTTCAAGGACGAC
GGCAACTACAAGACCCGCGCCGAGGTGAAGTTCGAGGGCGACACCCTGGTGAACCGCAT
CGAGCTGAAGGGCATCGACTTCAAGGAGGACGGCAACATCCTGGGGCACAAGCTGGAGTA
CAACTACAACAGCCACAACGTCTATATCACCGCCGACAAGCAGAAGAACGGCATCAAGGCA
AACTTCAAGATCCGCCACAACATCGAGGACGGCGGCGTGCAGCTCGCCGACCACTACCAG
CAGAACACCCCCATCGGCGACGGCCCCGTGCTGCTGCCCGACAACCACTACCTGAGCTAC
CAGTCCAAACTGAGCAAAGACCCCAACGAGAAGCGCGATCACATGGTCCTGCTGGAGTTC
GTGACCGCCGCCGGGATCACTCTCGGCATGGACGAGCTGTACAAGGGTTCTGGAAGTGGA
TCCACGATTGACCAGTGGCTGCTGAAAAACGCGAAAGAAGATGCTATTGCAGAACTGAAAA
AGGCTGGTATCACCGCTGACTTTTACTTCAACGTGATCAATAAAGCGAACTATGTGTGGTCT

GTTAACTTGTATAAGAACGATATCCTGAAAGCTCACGCCGGGAGCTCTGGACTCGAGCACCACCACCACCACCACTAA

**P22 NTOM20-mVenus-Ash0.8-His6**

ATGGTGGGCCGCAACAGCGCGATTGCAGCTGGAGTGTGCGGCGCGCTGTTCATTGGCTATTGCATTTACTTTGATCGCAAACGCCGCAGCGATCCGAACTTTAAATCTAGAGTGAGCAAGGGCGAGGAGCTGTTCACCGGGGTGGTGCCCATCCTGGTCGAGCTGGACGGCGACGTAAACGGCCACAAGTTCAGCGTGTCCGGCGAGGGCGAGGGCGATGCCACCTACGGCAAGCTGACCCTGAAGTTCATCAGCACCACCGGCAAGCTGCCCGTGCCCTGGCCCACCCTCGTGACCACCCTCGGCTACGGCCTGCAGTGCTTCGCCCGCTACCCCGACCACATGAAGCAGCACGACTTCTTCAAGTCCGCCATGCCCGAAGGCTACGTCCAGGAGCGCACCATCTTCTTCAAGGACGACGGCAACTACAAGACCCGCGCCGAGGTGAAGTTCGAGGGCGACACCCTGGTGAACCGCATCGAGCTGAAGGGCATCGACTTCAAGGAGGACGGCAACATCCTGGGGCACAAGCTGGAGTACAACTACAACAGCCACAACGTCTATATCACCGCCGACAAGCAGAAGAACGGCATCAAGGCAAACTTCAAGATCCGCCACAACATCGAGGACGGCGGCGTGCAGCTCGCCGACCACTACCAGCAGAACACCCCCATCGGCGACGGCCCCGTGCTGCTGCCCGACAACCACTACCTGAGCTACCAGTCCAAACTGAGCAAAGACCCCAACGAGAAGCGCGATCACATGGTCCTGCTGGAGTTCGTGACCGCCGCCGGGATCACTCTCGGCATGGACGAGCTGTACAAGGGTTCTGGAAGTGGATCCACGATTGACCAGTGGCTGCTGAAAAACGCGAAAGAAGATGCTATTGCAGAACTGAAAAAGGCTGGTATCACCTCTGACCGTGAATTCAACGCGATCAATAAAGCGAATTATGTGTGGTCTGTTAACGTGTACAAGAACGAGATCCTGAAAGCTCACGCCGGGAGCTCTGGACTCGAGCACCACCACCACCACCACTAA

**P23 NTOM20-mVenus-Ash0.9-His6**

ATGGTGGGCCGCAACAGCGCGATTGCAGCTGGAGTGTGCGGCGCGCTGTTCATTGGCTATTGCATTTACTTTGATCGCAAACGCCGCAGCGATCCGAACTTTAAATCTAGAGTGAGCAAGGGCGAGGAGCTGTTCACCGGGGTGGTGCCCATCCTGGTCGAGCTGGACGGCGACGTAAACGGCCACAAGTTCAGCGTGTCCGGCGAGGGCGAGGGCGATGCCACCTACGGCAAGCTGACCCTGAAGTTCATCAGCACCACCGGCAAGCTGCCCGTGCCCTGGCCCACCCTCGTGACCACCCTCGGCTACGGCCTGCAGTGCTTCGCCCGCTACCCCGACCACATGAAGCAGCACGACTTCTTCAAGTCCGCCATGCCCGAAGGCTACGTCCAGGAGCGCACCATCTTCTTCAAGGACGACGGCAACTACAAGACCCGCGCCGAGGTGAAGTTCGAGGGCGACACCCTGGTGAACCGCATCGAGCTGAAGGGCATCGACTTCAAGGAGGACGGCAACATCCTGGGGCACAAGCTGGAGTACAACTACAACAGCCACAACGTCTATATCACCGCCGACAAGCAGAAGAACGGCATCAAGGCAAACTTCAAGATCCGCCACAACATCGAGGACGGCGGCGTGCAGCTCGCCGACCACTACCAG

CAGAACACCCCCATCGGCGACGGCCCCGTGCTGCTGCCCGACAACCACTACCTGAGCTAC
CAGTCCAAACTGAGCAAAGACCCCAACGAGAAGCGCGATCACATGGTCCTGCTGGAGTTC
GTGACCGCCGCCGGGATCACTCTCGGCATGGACGAGCTGTACAAGGGTTCTGGAAGTGGA
TCCACGATTGACCAGTGGCTGCTGAAAAACGCGAAAGAAGATGCTATTGCAGAACTGAAAA
AGGCTGGTATCACCTCAGACCGGGACTTCAACGTGATCAATAAAGCGAATCATGTGTGGTCT
GTTAACTTGTATAAGAACCACATCCTGAAAGCTCACGCCGGGAGCTCTGGACTCGAGCACC
ACCACCACCACCACTAA

**P24 FUS$_N$-mCheery-Ash1**

ATGGCCTCAAACGATTATACCCAACAAGCAACCCAAAGCTATGGGGCCTACCCCACCCAGC
CCGGGCAGGGCTATTCCCAGCAGAGCAGTCAGCCCTACGGACAGCAGAGTTACAGTGGTT
ATAGCCAGTCCACGGACACTTCAGGCTATGGCCAGAGCAGCTATTCTTCTTATGGCCAGAG
CCAGAACACAGGCTATGGAACTCAGTCAACTCCCCAGGGATATGGCTCGACTGGCGGCTAT
GGCAGTAGCCAGAGCTCCCAATCGTCTTACGGGCAGCAGTCCTCCTATCCTGGCTATGGCC
AGCAGCCAGCTCCCAGCAGCACCTCGGGAAGTTACGGTAGCAGTTCTCAGAGCAGCAGCT
ATGGGCAGCCCCAGAGTGGGAGCTACAGCCAGCAGCCTAGCTATGGTGGACAGCAGCAAA
GCTATGGACAGCAGCAAAGCTATAATCCCCCTCAGGGCTATGGACAGCAGAACCAGTACAA
CAGCAGCAGTGGTGGTGGAGGTGGAGGTGGAGGTGGAGGTAACTATGGCCAAGATCAATC
CTCCATGAGTAGTGGTGGTGGCAGTGGTGGCGGTTATGGCAATCAAGACCAGAGTGGTGG
AGGTGGCAGCGGTGGCTATGGACAGCAGGACCGTGGAGGTGGAGGAATGGTGTCTAAAG
GCGAGGAGGATAACATGGCTATAATCAAGGAATTTATGAGGTTCAAGGTGCATATGGAGGGA
TCTGTGAACGGTCACGAGTTCGAAATCGAGGGCGAAGGCGAGGGGCGACCCTATGAAGG
GACACAGACGGCCAAACTGAAGGTGACCAAGGGTGGCCCCCTGCCTTTCGCCTGGGACAT
CCTGTCACCCCAGTTTATGTACGGCTCTAAGGCTTACGTTAAGCACCCTGCCGATATTCCGG
ACTACTTGAAACTGAGCTTCCCTGAAGGGTTCAAGTGGAACGGGTAATGAATTTTGAGGAT
GGCGGTGTAGTCACAGTTACCCAGGACAGCTCTCTTCAGGACGGAGAATTTATCTATAAGGT
TAAACTTCGGGGCACTAACTTCCCATCAGACGGCCCCGTGATGCAGAAGAAAACTATGGGC
TGGGAAGCCAGCTCAGAACGCATGTATCCCGAGGACGGAGCACTGAAGGGAGAAATCAAG
CAGCGACTCAAGCTGAAGGATGGGGGACATTATGATGCTGAGGTCAAAACCACCTATAAGG
CCAAGAAACCCGTTCAGCTTCCTGGGGCATACAACGTGAATATCAAACTGGATATTACATCC
CATAACGAAGACTATACCATCGTGGAACAGTACGAGCGGGCCGAGGGCAGACATAGCACAG
GGGGCATGGATGAATTGTACAAGGCGGCCGGATCCGGTGGTACCTCTGGAGGTACGATTG
ACCAGTGGCTGCTGAAAAACGCGAAAGAAGATGCTATTGCAGAACTGAAAAAGGCTGGTAT

CACCGCTGACTTTTACTTCAACGTGATCAATAAAGCGAACTATGTGTGGTCTGTTAACTTGTA
TAAGAACGATATCCTGAAAGCTCACGCCTGA

**P25 NLS-FenixS-EGFP-FTH1**

ATGGCGGAGCCGCGTAGCAAACGTCCGCGTGTGACGCTGCCGGCGTGTCCGGCGCATAG
TGGGGAGTTTGGTTCTGGCTCAATGTACCAGGACAAGATGTTGAATGACACAGTAGCAAAG
GTGCGACAGATCCTGCAGGCGGACAGGGTGGTTATGTTCCAATTCGAGGAGGATTACAGC
GGAGAGGTGGTGGTGGAGGCCGTGGACGGTAGGTGGAGCTCCATCCTGGGGACCCAGTG
TAGTGATAGTTACTTCATGGAGTCAAGGGGCGAGGAGTATGCTCGCGGCCGCTACCAGGCC
ATCGCCGACATCTACACCGCTAACCTGTCAGAGTGCCACAGGGATCTGCTGGCGCAGTTTC
AGGTGAGAGCGGTCCTGGCCGTGCCCATCCTGCAGGGCAAGAAGCTGTGGGGCCTGTTG
GTGGCACACCAGCTGGCGGCCCCTAGACAGTGGCAGTCCTGGGAGATCGACTTTCTGAAG
CAGCAGGCCGTGGCGGTGGGCATAGCCATCCAACAGATCGGCGGCTCCGGCTCTGGATC
CGGAGCTACCATGAGTAAAGGAGAAGAACTTTTCACTGGAGTTGTCCCAATTCTTGTTGAAT
TAGATGGTGATGTTAATGGGCACAAATTTTCTGTCAGTGGAGAGGGTGAAGGTGATGCAACA
TACGGAAAACTTACCCTTAAATTTATTTGCACTACTGGAAAACTACCTGTTCCATGGCCAACA
CTTGTCACTACCCTGACTTATGGTGTTCAATGCTTTTCAAGATACCCAGATCATATGAAACGG
CATGACTTTTTCAAGAGTGCCATGCCCGAAGGTTATGTACAGGAAAGAACTATATTTTTCAAA
GATGACGGGAACTACAAGACACGTGCTGAAGTCAAGTTTGAAGGTGATACCCTTGTTAATAG
AATCGAGTTAAAAGGTATTGATTTTAAAGAAGATGGAAACATTCTTGGACACAAATTGGAATA
CAACTATAACTCACACAATGTATACATCATGGCAGACAAACAAAAGAATGGAATCAAAGTTAA
CTTCAAAATTAGACACAACATTGAAGATGGAAGCGTTCAACTAGCAGACCATTATCAACAAAA
TACTCCAATTGGCGATGGCCCTGTCCTTTTACCAGACAACCATTACCTGTCCACACAATCTAA
GCTTTCGAAAGATCCCAACGAAAAGAGAGACCACATGGTCCTTCTTGAGTTTGTAACAGCT
GCTGGGATTACACATGGCATGGATGAACTATACAAAGGTGGATCTGGAGGTTCAGGTGGAA
GTGCGGCCGGTACCGGCAGCGGAGGAATGACGACCGCGTCCACCTCGCAGGTGCGCCAG
AACTACCACCAGGACTCAGAGGCCGCCATCAACCGCCAGATCAACCTGGAGCTCTACGCC
TCCTACGTTTACCTGTCCATGTCTTACTACTTTGACCGCGATGATGTGGCTTTGAAGAACTTT
GCCAAATACTTTCTTCACCAATCTCATGAGGAGAGGGAACATGCTGAGAAACTGATGAAGCT
GCAGAACCAACGAGGTGGCCGAATCTTCCTTCAGGATATCAAGAAACCAGACTGTGATGAC
TGGGAGAGCGGGCTGAATGCAATGGAGTGTGCATTACATTTGGAAAAAAATGTGAATCAGT
CACTACTGGAACTGCACAAACTGGCCACTGACAAAAATGACCCCCATTTGTGTGACTTCATT
GAGACACATTACCTGAATGAGCAGGTGAAAGCCATCAAAGAATTGGGTGACCACGTGACCA

ACTTGCGCAAGATGGGAGCGCCCGAATCTGGCTTGGCGGAATATCTCTTTGACAAGCACAC
CCTGGGAGACAGTGATAATGAAAGCTGA

**P30 FenixS-VP16-NLS-IRES-E-Ash1**

ATGTACCAGGACAAGATGTTGAATGACACAGTAGCAAAGGTGCGACAGATCCTGCAGGCGG
ACAGGGTGGTTATGTTCCAATTCGAGGAGGATTACAGCGGAGAGGTGGTGGTGGAGGCCG
TGGACGGTAGGTGGAGCTCCATCCTGGGGACCCAGTGTAGTGATAGTTACTTCATGGAGTC
AAGGGGCGAGGAGTATGCTCGCGGTCGCTACCAGGCCATCGCCGACATCTACACCGCTAA
CCTGTCAGAGTGCCACAGGGATCTGCTGGCGCAGTTTCAGGTGAGAGCGGTCCTGGCCGT
GCCCATCCTGCAGGGCAAGAAGCTGTGGGGCCTGTTGGTGGCACACCAGCTGGCGGCCC
CTAGACAGTGGCAGTCCTGGGAGATCGACTTTCTGAAGCAGCAGGCCGTGGCGGTGGGC
ATAGCCATCCAACAGATCGAATTCGATAGTGCTGGTAGTGCTGGTAGTGCTGGTTCCGCGTA
CAGCCGCGCGCGTACGAAAAACAATTACGGGTCTACCATCGAGGGCCTGCTCGATCTCCC
GGACGACGACGCCCCCGAAGAGGCGGGGCTGGCGGCTCCGCGCCTGTCCTTTCTCCCC
GCGGGACACACGCGCAGACTGTCGACGGCCCCCCCGACCGATGTCAGCCTGGGGGACGA
GCTCCACTTAGACGGCGAGGACGTGGCGATGGCGCATGCCGACGCGCTAGACGATTTCGA
TCTGGACATGTTGGGGGACGGGGATTCCCCGGGTCCGGGATTTACCCCCCACGACTCCGC
CCCCTACGGCGCTCTGGATATGGCCGACTTCGAGTTTGAGCAGATGTTTACCGATGCCCTT
GGAATTGACGAGTACGGTGGGCCCAAGAAAAAGCGGAAGGTGTGATCTAGAGTCGACCTG
CAGCCCAAGCTTAAAACAGCTCTGGGGTTGTACCCACCCCAGAGGCCCACGTGGCGGCTA
GTACTCCGGTATTGCGGTACCCTTGTACGCCTGTTTTATACTCCCTTCCCGTAACTTAGACGC
ACAAAACCAAGTTCAATAGAAGGGGGTACAAACCAGTACCACCACGAACAAGCACTTCTGT
TTCCCCGGTGATGTCGTATAGACTGCTTGCGTGGTTGAAAGCGACGGATCCGTTATCCGCT
TATGTACTTCGAGAAGCCCAGTACCACCTCGGAATCTTCGATGCGTTGCGCTCAGCACTCA
ACCCCAGAGTGTAGCTTAGGCTGATGAGTCTGGACATCCCTCACCGGTGACGGTGGTCCA
GGCTGCGTTGGCGGCCTACCTATGGCTAACGCCATGGGACGCTAGTTGTGAACAAGGTGT
GAAGAGCCTATTGAGCTACATAAGAATCCTCCGGCCCCTGAATGCGGCTAATCCCAACCTC
GGAGCAGGTGGTCACAAACCAGTGATTGGCCTGTCGTAACGCGCAAGTCCGTGGCGGAAC
CGACTACTTTGGGTGTCCGTGTTTCCTTTTATTTTATTGTGGCTGCTTATGGTGACAATCACA
GATTGTTATCATAAAGCGAATTGGATTGCGGCGATATCGCCACCATGCCCCGCCCCAAGCTC
AAGTCCGATGACGAGGTACTCGAGGCCGCCACCGTAGTGCTGAAGCGTTGCGGTCCCATA
GAGTTCACGCTCAGCGGAGTAGCAAAGGAGGTGGGGCTCTCCCGCGCAGCGTTAATCCAG
CGCTTCACCAACCGCGATACGCTGCTGGTGAGGATGATGGAGCGCGGCGTCGAGCAGGT
GCGGCATTACCTGAATGCGATACCGATAGGCGCAGGGCCGCAAGGGCTCTGGGAATTTTG

CAGGTGCTCGTTCGGAGCATGAACACTCGCAACGACTTCTCGGTGAACTATCTCATCTCCT
GGTACGAGCTCCAGGTGCCGGAGCTACGCACGCTTGCGATCCAGCGGAACCGCGCGGTG
GTGGAGGGGATCCGCAAGCGACTGCCCCCAGGTGCTCCTGCGGCAGCTGAGTTGCTCCT
GCACTCGGTCATCGCTGGCGCGACGATGCAGTGGGCCGTCGATCCGGATGGTGAGCTAGC
TGATCATGTGCTGGCTCAGATCGCTGCCATCCTGTGTTTAATGTTTCCCGAACACGACGATT
TCCAACTCCTCCAGGCACATGCGTCCGCGTACAGCTTAATTGTACATAACAGTGCTGGTAGT
GCTGGTAGTGCTGGTACGATTGACCAGTGGCTGCTGAAAAACGCGAAAGAAGATGCTATTG
CAGAACTGAAAAAGGCTGGTATCACCGCTGACTTTTACTTCAACGTGATCAATAAAGCGAAC
TATGTGTGGTCTGTTAACTTGTATAAGAACGATATCCTGAAAGCTCACGCCTACCCATACGAT
GTTCCAGATTACGCTTAA

**P31 FenixS-FUS-VP16-NLS-IRES-E-Ash1**

ATGTACCAGGACAAGATGTTGAATGACACAGTAGCAAAGGTGCGACAGATCCTGCAGGCGG
ACAGGGTGGTTATGTTCCAATTCGAGGAGGATTACAGCGGAGAGGTGGTGGTGGAGGCCG
TGGACGGTAGGTGGAGCTCCATCCTGGGGACCCAGTGTAGTGATAGTTACTTCATGGAGTC
AAGGGGCGAGGAGTATGCTCGCGGTCGCTACCAGGCCATCGCCGACATCTACACCGCTAA
CCTGTCAGAGTGCCACAGGGATCTGCTGGCGCAGTTTCAGGTGAGAGCGGTCCTGGCCGT
GCCCATCCTGCAGGGCAAGAAGCTGTGGGGCCTGTTGGTGGCACACCAGCTGGCGGCCC
CTAGACAGTGGCAGTCCTGGGAGATCGACTTTCTGAAGCAGCAGGCCGTGGCGGTGGGC
ATAGCCATCCAACAGATCGAATTCATGGCCTCAAACGATTATACCCAACAAGCAACCCAAAG
CTATGGGGCCTACCCCACCCAGCCCGGGCAGGGCTATTCCCAGCAGAGCAGTCAGCCCTA
CGGACAGCAGAGTTACAGTGGTTATAGCCAGTCCACGGACACTTCAGGCTATGGCCAGAGC
AGCTATTCTTCTTATGGCCAGAGCCAGAACACAGGCTATGGAACTCAGTCAACTCCCCAGG
GATATGGCTCGACTGGCGGCTATGGCAGTAGCCAGAGCTCCCAATCGTCTTACGGGCAGCA
GTCCTCCTACCCTGGCTATGGCCAGCAGCCAGCTCCCAGCAGCACCTCGGGAAGTTACGG
TAGCAGTTCTCAGAGCAGCAGCTATGGGCAGCCCCAGAGTGGGAGCTACAGCCAACAGCC
TAGCTATGGTGGACAGCAGCAATCTTACGGTCAACAACAGAGCTATAATCCCCCTCAGGGCT
ATGGACAGCAGAACCAGTACAACAGCAGCAGTGGTGGTGGAGGTGGAGGTGGAGGTGGA
GGTAACTATGGCCAAGATCAATCCTCCATGAGTAGTGGTGGTGGCAGTGGTGGCGGTTATG
GCAATCAAGACCAGAGTGGTGGAGGTGGCAGCGGTGGCTATGGACAGCAGGACCGTGGA
GCGCGTACGAAAAACAATTACGGGTCTACCATCGAGGGCCTGCTCGATCTCCCGGACGAC
GACGCCCCCGAAGAGGCGGGGCTGGCGGCTCCGCGCCTGTCCTTTCTCCCCGCGGAC
ACACGCGCAGACTGTCGACGGCCCCCCCGACCGATGTCAGCCTGGGGGACGAGCTCCAC
TTAGACGGCGAGGACGTGGCGATGGCGCATGCCGACGCGCTAGACGATTTCGATCTGGAC

ATGTTGGGGGACGGGGATTCCCCGGGTCCGGGATTTACCCCCCACGACTCCGCCCCCTAC
GGCGCTCTGGATATGGCCGACTTCGAGTTTGAGCAGATGTTTACCGATGCCCTTGGAATTG
ACGAGTACGGTGGGCCCAAGAAAAAGCGGAAGGTGTGAATCTAGAGTCGACCTGCAGCCCA
AGCTTAAAACAGCTCTGGGGTTGTACCCACCCCAGAGGCCCACGTGGCGGCTAGTACTCC
GGTATTGCGGTACCCTTGTACGCCTGTTTTATACTCCCTTCCCGTAACTTAGACGCACAAAA
CCAAGTTCAATAGAAGGGGGTACAAACCAGTACCACCACGAACAAGCACTTCTGTTTCCCC
GGTGATGTCGTATAGACTGCTTGCGTGGTTGAAAGCGACGGATCCGTTATCCGCTTATGTAC
TTCGAGAAGCCCAGTACCACCTCGGAATCTTCGATGCGTTGCGCTCAGCACTCAACCCCAG
AGTGTAGCTTAGGCTGATGAGTCTGGACATCCCTCACCGGTGACGGTGGTCCAGGCTGCG
TTGGCGGCCTACCTATGGCTAACGCCATGGGACGCTAGTTGTGAACAAGGTGTGAAGAGCC
TATTGAGCTACATAAGAATCCTCCGGCCCCTGAATGCGGCTAATCCCAACCTCGGAGCAGG
TGGTCACAAACCAGTGATTGGCCTGTCGTAACGCGCAAGTCCGTGGCGGAACCGACTACT
TTGGGTGTCCGTGTTTCCTTTTATTTTATTGTGGCTGCTTATGGTGACAATCACAGATTGTTAT
CATAAAGCGAATTGGATTGCGGCGATATCGCCACCATGCCCCGCCCCAAGCTCAAGTCCGA
TGACGAGGTACTCGAGGCCGCCACCGTAGTGCTGAAGCGTTGCGGTCCCATAGAGTTCAC
GCTCAGCGGAGTAGCAAAGGAGGTGGGGCTCTCCCGCGCAGCGTTAATCCAGCGCTTCAC
CAACCGCGATACGCTGCTGGTGAGGATGATGGAGCGCGGCGTCGAGCAGGTGCGGCATTA
CCTGAATGCGATACCGATAGGCGCAGGGCCGCAAGGGCTCTGGGAATTTTTGCAGGTGCT
CGTTCGGAGCATGAACACTCGCAACGACTTCTCGGTGAACTATCTCATCTCCTGGTACGAG
CTCCAGGTGCCGGAGCTACGCACGCTTGCGATCCAGCGGAACCGCGCGGTGGTGGAGGG
GATCCGCAAGCGACTGCCCCCAGGTGCTCCTGCGGCAGCTGAGTTGCTCCTGCACTCGGT
CATCGCTGGCGCGACGATGCAGTGGGCCGTCGATCCGGATGGTGAGCTAGCTGATCATGT
GCTGGCTCAGATCGCTGCCATCCTGTGTTTAATGTTTCCCGAACACGACGATTTCCAACTCC
TCCAGGCACATGCGTCCGCGTACAGCTTAATTGTACATAACAGTGCTGGTAGTGCTGGTAGT
GCTGGTACGATTGACCAGTGGCTGCTGAAAAACGCGAAAGAAGATGCTATTGCAGAACTGA
AAAAGGCTGGTATCACCGCTGACTTTTACTTCAACGTGATCAATAAAGCGAACTATGTGTGG
TCTGTTAACTTGTATAAGAACGATATCCTGAAAGCTCACGCCTACCCATACGATGTTCCAGAT
TACGCTTAA
